# Correction of amyotrophic lateral sclerosis related phenotypes in induced pluripotent stem cell-derived motor neurons carrying a hexanucleotide expansion mutation in *C9orf72* by CRISPR/Cas9 genome editing using homology-directed repair

**DOI:** 10.1101/2019.12.17.864520

**Authors:** Nidaa Ababneh, Jakub Scaber, Rowan Flynn, Andrew Douglas, Martin R. Turner, David Sims, Ruxandra Dafinca, Sally A. Cowley, Kevin Talbot

## Abstract

The G4C2 hexanucleotide repeat expansion (HRE) in *C9orf72* is the commonest cause of familial amyotrophic lateral sclerosis (ALS). A number of different methods have been used to generate isogenic control lines using CRISPR (clustered regularly interspaced short palindromic repeats)/Cas9 and non-homologous end-joining (NHEJ) by deleting the repeat region with the risk of creating indels and genomic instability. In this study we demonstrate complete correction of an induced pluripotent stem cell (iPSC) line derived from a *C9orf72*-HRE positive ALS/FTD patient using CRISPR/Cas9 genome editing and homology directed repair (HDR), resulting in replacement of the excised region with a donor template carrying the wild-type repeat size to maintain the genetic architecture of the locus. The isogenic correction of the *C9orf72* HRE restored normal expression and methylation at the C9orf72 locus, reduced intron retention in the edited lines, and abolished pathological phenotypes associated with the *C9orf72* HRE expansion in iPSC derived motor neurons (iPSMNs).

RNA sequencing of the mutant line identified 2220 differentially expressed genes compared to its isogenic control. Enrichment analysis demonstrated an over-representation of ALS relevant pathways, including calcium ion dependent exocytosis, synaptic transport and the KEGG ALS pathway, as well as new targets of potential relevance to ALS pathophysiology.

Complete correction of the *C9orf72* HRE in iPSMNs by CRISPR/Cas9 mediated HDR provides an ideal model to study the earliest effects of the hexanucleotide expansion on cellular homeostasis and the key pathways implicated in ALS pathophysiology.

## Introduction

Amyotrophic lateral sclerosis (ALS) is a rapidly progressive and uniformly fatal adult-onset neurodegenerative disorder characterized by loss of motor neurons (MNs) in the brain and spinal cord, with an average survival of approximately 2.5 years from symptom onset [1]. The clinical, genetic and pathological overlap with frontotemporal dementia (FTD) is now well established [2]. A hexanucleotide repeat expansion (HRE) mutation in the *C9orf72* gene is responsible for 35-40% of cases of familial ALS (fALS), 5-7% of sporadic ALS (sALS), and approximately 25% of cases of familial frontotemporal dementia (fFTD) in populations of Northern European genetic heritage [3,4]. The hexanucleotide expansion is located in a non-coding region of the gene which forms either the upstream promoter or the first intron of *C9orf72* transcripts. The number of (G_4_C_2_)_n_ hexanucleotide repeats in affected individuals is typically more than 1000 [5,6].

Multiple mechanisms of *C9orf72*-HRE toxicity have been proposed. The accumulation of (G4C2)_n_ transcripts in RNA foci in neuronal nuclei may lead to the sequestration of RNA binding proteins, with disruption of RNA handling or induction of nucleolar stress. Repeat-associated non-ATG (RAN) translation across the G4C2 repeat expansion in both sense and antisense directions produces aggregation-prone, potentially toxic, poly-dipeptide repeats (DPRs) [7,8]. In addition, hexanucleotide repeat expansions result in transcriptional downregulation of *C9orf72* mRNA and reduced protein levels in affected regions of the brain of ALS/FTD patients [9], suggesting that haploinsufficiency may also contribute to the phenotype.

RNA foci and RAN-translation products can be detected in induced pluripotent stem cell (iPSC)-derived motor neurons (iPSMNs) derived from patients with *C9orf72* hexanucleotide expansions, in association with varied phenotypes, including altered neuronal excitability, sequestration of RNA-binding proteins, increased endoplasmic reticulum (ER) stress and defects in nucleocytoplasmic transport and autophagy [10–16].

A key challenge in using techniques such as RNA sequencing for the identification of disease mechanisms using iPSC neuronal models is that inherent biological variance arising from differences in genetic background, clonal selection, and inconsistencies in reprogramming and differentiation protocols, may be greater than the expression changes due to a disease-causing mutation. Genome editing techniques, such as CRISPR (clustered regularly interspaced short palindromic repeats)/Cas9 and zinc-finger nuclease mediated-gene targeting, have been employed to remove point mutations in iPSCs from an ALS patients with SOD1 mutations and in iPSCs from an FTD patient with a progranulin mutation, to create isogenic control lines [17–20]. Recently, the successful excision of *C9orf72* (G4C2)_n_ using CRISPR/Cas9 technology has also been reported, yielding isogenic lines with a missing repeat region [16].

Here, we report the generation of isogenic iPSC line from an ALS/FTD patient carrying an expansion of 1000 G4C2 repeats in *C9orf72* using CRISPR/Cas9-mediated HDR to replace the pathogenic expansion with a donor template carrying a normal repeat size of (G4C2)_2_. To reduce the possibility of off-target mutations and improve on-target specificity, a double-nicking approach with the D10A mutant nickase version of Cas9 (Cas9n) was used [21]. In MNs differentiated from the corrected iPSC lines, we demonstrate the correction of pathological markers of the *C9orf72* HRE, such as sense and antisense RNA foci and dipeptide toxicity. We also demonstrate restoration of *C9orf72* transcript expression and methylation to levels seen in normal controls. Using RNA sequencing, we identify dysregulation of ALS relevant pathways specific to lines carrying the expansion, including glutamate excitotoxicity and cell death and the functional validation of selected pathways in the corrected cell lines. This study therefore demonstrates that HDR-mediated correction in patient-derived iPSCs can lead to the generation of induced MN-like cells that display features characteristic of normal cells, which has the advantage over other techniques of generating a more precise isogenic control line free of indels.

## Materials and Methods

### Fibroblast reprogramming

Skin fibroblasts obtained from an ALS/FTD patient (C9-02) carrying the (G4C2)_n_ repeat expansion mutation in the *C9orf72* gene, confirmed by repeat primed PCR and Southern blotting, were used to generate two iPSC lines (C9-02-02 & C0-02-03) using the CytoTune-1-iPSC reprogramming kit (Life Technologies), as previously described [10]. Ethical approval for the collection and use of these cells was obtained from the South Wales Research Ethics Committee (approval number: 12/WA/0186).

### Karyotyping

Genomic DNA was extracted using the DNeasy kit (QIAGEN). Genome integrity for iPSC lines and their corresponding fibroblasts was assessed using the Human OmniExpress24 array (∼700,000 markers, Illumina) and analysed using KaryoStudio software (Illumina).

### CRISPR/Cas9 gRNA construct preparation and guide RNA construction

Guide-RNAs (gRNA1 and gRNA2) were designed to target a sequence 140 bps upstream of the repeat region. All PCR reactions were done using Phusion High Fidelity DNA Polymerase (Thermo Fisher Scientific) in a 50 μl total volume. Sanger sequencing was performed by Source BioScience (Oxford, United Kingdom). Oligo sequences for gRNA synthesis are listed in Supplementary Table 2. gRNA synthesis was performed using the pX335 vector (Addgene, Plasmid #42335) containing a D10A nickase for the co-expression of SpCas9 and gRNA, following a published protocol [22]. For the construction of gRNA 1&2, the pX335 vector was linearized with BbsI and gel-purified. The two oligos for each gRNA were annealed, phosphorylated, and ligated into the pX335 vector. The vector was then transformed into Stabl2 E Coli and DNA minipreps were prepared and sequenced to check for the presence of correct gRNA sequences. Finally, two midipreps were prepared for successfully constructed gRNA 1 & 2.

### Design of homology-directed repair template

The DNA regions upstream and downstream of the repeat were assessed individually for the presence of any single nucleotide polymorphisms (SNPs) using two sets of specific primers listed in Supplementary Table 3. The left and right homology arms were amplified from the wild-type allele of the same patient DNA (C9-02), which contains two G4C2 repeats. The amplification was performed using two units of Phusion polymerase with 10X buffer (NEB), 10 μM forward primer, 10 μM reverse primer, 10 mM dNTP mix and 5 ng DNA (HR-F and HR-R primers in Supplementary Table 3). The mixture was heated to 98°C for 2 minutes, then 35 cycles of 98°C for 15 seconds, 67°C for 20 seconds and 72°C for 3 minutes and followed by 5 minutes of final extension at the same temperature. The purified PCR product was inserted into the pGEMT easy vector (Promega) according to the manufacturer’s instructions. Two Nsi1 sites upstream and downstream of the repeat region were inserted using site-directed mutagenesis (SDM) to allow the evaluation of successfully targeted clones by polymerase chain reaction-restriction fragment length polymorphism (PCR-RFLP), using the same PCR reaction conditions listed above (SDM-F and SDM-R primers in Supplementary table 3). After that, 1 μl of DpnI restriction enzyme was directly added to the PCR product and incubated for 30 minutes at 37°C to remove the original plasmid. A LoxP-flanked EF1α-Puro-Tk cassette in a pENTR plasmid (Invitrogen) was inserted between the two homology arms ∼100bp upstream of the repeat region and in between the two Nsi1 sites. The cassette was amplified by PCR then ligated to the donor template using Gibson Assembly Master Mix (NEB, primers in Supplementary Table 3). The reaction was incubated at 50 °C for 15 minutes, followed by bacterial transformation and miniprep preparation. The resulting construct was checked by direct sequencing and midiprep was prepared.

### Cell Culture and Transfection of HEK293T cells for cleavage activity testing

Genomic DNA samples from untransfected and transfected cells were extracted using DNeasy Blood & Tissue kit (Qiagen). The genomic region flanking the CRISPR cleavage site was amplified by PCR using specific Surv-F and Surv-R primers to amplify 900 bp surrounding the genomic cleavage site (Supplementary Table 3) and the resulting products were purified using PCR Cleanup kit (Qiagen). Cleavage assays were performed using 400ng of PCR products that were denatured by heating at 95°C then reannealed to form heteroduplex DNA using a thermocycler. Then, the reannealed PCR products were digested with T7 endonuclease 1 (T7E1, New England Biolabs). Digestion was performed using 10 unites of T7E1 enzyme and incubated for 15 minutes at 37°C. The T7EI reaction was stopped by adding 2 μl of 0.25M EDTA solution. The digested products were analysed on 3% agarose gel electrophoresis stained with Ethidium Bromide to check for heteroduplex DNA formation.

### CRISPR-Mediated genome targeting of iPSCs

On day 0, iPSCs were pre-treated with 2 μM of Rho Kinase (ROCK) inhibitor for at least 1 h prior to nucleofection. Cells were dissociated into single cells by incubating them with TrypLE for 5 minutes at 37°C, mixing with PBS and followed by centrifugation at 400g for 5 minutes. Two million cells were transfected with 2.5 μg of each gRNA-encoding plasmid and 4 μg of donor plasmid using the Neon Transfection system (Life Technologies). A GFP puromycin-free plasmid was used as a positive control. Following electroporation, cells were plated on 10 cm dishes coated with drug-resistant (DR4) mouse embryonic fibroblasts (MEFs) in human embryonic stem cell (hES) media composed of Knock-out Dulbecco’s modified Eagle’s medium (DMEM) (Gibco), 10% knock-out-serum replacement (Gibco), 2 mM Glutamax-I (Gibco) 100 U/mL, Penicillin/Streptomycin (P/S) (Gibco), 1% Nonessential amino acids (NEAA) (Gibco), 0.5 mM 2-Mercaptoethanol (Gibco), and 10 ng/mL Basic fibroblast growth factor (bFGF) (R&D) and also supplemented with 10 mM ROCK inhibitor Y-27632 (RI) for overnight. On day 3, puromycin was added at 0.35 μg/mL to initiate the selection process. Surviving individual colonies were picked and expanded in 24-well plates coated with DR4 MEFs. Confluent wells were further split for expansion and molecular screening.

### Repeat-Primed PCR

iPSC clones were grown and passaged on two 24-well plates for DNA extraction and freezing in LN for storage. Total DNA was extracted using QIAamp DNA Blood Mini Kit (QIAGEN) and resuspended in Nuclease free water. The clones were screened first by Repeat-Primed PCR (RP-PCR). Primer sequences used in RP-PCR are listed in Supplementary Table 3. Two RP-PCRs were performed, RP-PCR1 to assess the presence of the repeats on the 5’ direction, and RP-PCR2 to assess the presence of the repeats on the 3’ direction. RP-PCR assays were performed as previously described with a brief modification on the primer sequences [9,23]. Both PCR mixes were prepared individually in a reaction volume of 15 µl, containing 100ng of genomic DNA, 8 µl Extensor mastermix (Thermo Scientific), 6 µl of 5 M Betaine, 20 µM of both forward and repeat specific primer and 2 µM of reverse primer. The reactions were subjected to touchdown PCR programme consisting of 94°C for 30 seconds, 60°C for 30 seconds and 72°C for 30 seconds, then 1 µl of each PCR product was mixed with 8 µl of Formamide, and 0.5 µl GeneScan size standard (Applied Biosystems).

### PCR and direct sequencing of the repeat region

Following RP-PCR, 24 clones without the repeat expansion were screened by PCR amplification of the repeat region in a 50 µl reaction volume using Repeats-F and Repeats-R in Supplementary table 3. PCR was performed as follows: 2 minutes at 98°C, 35 cycles of 15 seconds at 98°C, 30 seconds at 67°C 2 minutes at 72°C and one final step for 5 minutes at 72°C. The PCR product was purified using QIAquick PCR Purification kit (QIAGEN). Sequencing analysis was performed using the Repeats-R primer (Source Bioscience company service (Oxford, UK).

### In-Out PCR and direct sequencing of intended modification

Clones without repeat expansion were screened by PCR amplification inside the EF1α-Puro-TK cassette and outside either 5’ or 3’ homology arms (5’-In-out, 3’In-out PCRs). Sequences and specifications of the primers are shown in Supplementary Table 3. PCR cycling conditions were performed as follows: 2 minutes at 98°C, 35 cycles of 15 seconds at 98°C, 30 seconds at 67°C and 2 minutes at 72°C and one final step for 5 minutes at 72°C. Thirty-five microliters of PCR products were purified using QIAquick PCR Purification kit (QIAGEN) and sequenced to confirm site-specific integration and successful insertion of Nsi1 (primers in Supplementary Table 3). Ten microliters of PCR products were digested with Nsi1 (3 hours with 5 units of Nsi1 enzyme (NEB)) and then, PCR fragments were visualised on 2% agarose gel electrophoresis to confirm the insertion of Nsi1 sites.

### Removal of Puro/Tk cassette using Cre-Lox P system

To excise the selection cassette, iPSCs from one of the heterozygous targeted clones (Ed-02) and one homozygous clone (Ed-03) were nucleofected using *Cre* mRNA following the same nucleofection experiment as described above. iPSCs were nucleofected with 0.5 μg *Cre* mRNA and seeded on 10 cm plates pre-coated with MEFs and maintained in hES supplemented with 10 μM Y-26732 and no antibiotics. After few days of nucleofection, cells were treated with 3 μg/ml Ganciclovir added directly to the feeding media. After 7-10 days, some of the growing colonies were picked up and transferred into 96-well plate coated with MEFs then split into two new plates, one used for freezing and one for genotyping and PCR analysis of Puro/Tk cassette-free clones. Multiple edited isogenic lines were generated from Ed-02 heterozygous clone after excision of the selection marker. Ed-03 derived clones were used as controls for Cre excision efficiency. Two edited clones were used after excision of the cassette Ed-02-01 and Ed-02-02 for phenotyping experiments. DNA was extracted from iPSCs of the Cre-treated and untreated edited clones and from the parental cells (C9-02-02). After that, PCR amplification was performed using 5’-Out-F and Puro-R primers to amplify inside the Puro/Tk cassette and outside the 5’-homology arm. Another PCR was carried out using Exc-F and Exc-R primers to amplify the repeat region, generating a fragment of 75 bp and 109 bp for the wild-type and the targeted allele, respectively. The difference in size is related to the presence of a LoxP site after excision of the cassette using Cre-LoxP system. All primers sequences are in Supplementary Table 3.

### Southern Blot

Southern blotting was applied on genomic DNA derived from edited cells, healthy controls and diseased cells. Briefly, DNA was extracted with DNeasy Blood & Tissue kit and 5-10 μg of genomic DNA was digested overnight at 37°C with *Alu*I- and *Dde*I. Then digested samples were separated by electrophoresis on 0.8% agarose gel, and then transferred overnight to a positively charged nylon membrane by capillary blotting (Roche Applied Science). In the next day, membrane was cross-linked by UV irradiation, and hybridized overnight at 55°C with 1 ng/ml digoxigenin-labelled (GGGGCC)^5^ oligonucleotide probe (Integrated DNA Technologies) in 25 ml hybridization reaction (EasyHyb Granules, Roche). Following denaturation of the probe at 95°C for 5 min, probe was snap cooled on ice for 30 seconds and immediately added to the prehybridised membrane. The following day, the membrane was washed twice with stringency washes (2X Saline-Sodium Citrate buffer (SSC) and 0.1% sodium dodecyl sulfate (SDS), each for 15 minutes at 65°C in the hybridisation oven. The membrane was then washed twice at the same temperature and conditions with 1X SSC and 0.1% SDS and finally washed twice with 0.5X SSC and 0.1% SDS for further 15 minutes at room temperature. The membrane was washed with 1X wash buffer (Roche Applied Science) for 2 minutes, then incubated with blocking solution (Roche Applied Science) for 1–2 hours. After that, the membrane was incubated with Anti-DIG-AP Fab fragments (Roche) for 25-30 minutes and washed three times with 1X washing buffer. Ready-to-use CDPD (Roche Applied Science) chemiluminescent substrate was added to the membrane and signals were visualised on Detection Film. All samples were compared to DIG-labelled DNA molecular-weight markers II (Roche Applied Science) and repeat number was estimated compared to the DNA marker sizes.

### Differentiation of iPSCs to motor neurons

iPSCs were differentiated to mature motor neurons (iPSMNs) using a previously described protocol [10], which was adapted with minor modification from the protocol described by Maury et. al [24]. Briefly, the iPSCs were grown on Matrigel until confluency, then neural induction was started using DMEM/F12:Neurobasal 1:1, N2, B27, ascorbic acid (0.5 mM), 2-mercaptoethanol, compound C (3 µM) and Chir99021 (3 µM). After 4 days in culture, RA (1 µM) and SAG (500 nM) were added to the medium. On the next day, Chir99021 and compound C were removed from the medium, and the cells were cultured for another 4–5 days. After that, cells were split 1:3 on approximately day 10. Subsequently, the medium was supplemented with growth factors BDNF (10 ng/mL), GDNF (10 ng/mL), N-[N-(3,5-Difluorophenacetyl)-L-ala-nyl]-S-phenylglycine t-butyl ester (DAPT) (10 mM), and laminin (0.5 mg/mL). After 7 days, DAPT and laminin were removed from the medium, and the neurons were allowed to mature until day 25-30.

### Immunofluorescence of motor neuron cultures

Cells on coverslips were washed once with PBS, then fixed with 4% paraformaldehyde for 15 minutes, permeabilized with 0.1% Triton X-100 in PBS and blocked in blocking buffer (0.01% TritonX-100/TBS with 10% normal goat serum) for one hour. Cells were subsequently incubated overnight at 4°C with primary antibodies: goat anti-ChAT, mouse anti-SMI-32, rabbit anti-Islet1/2 and rabbit anti-HB9, and goat anti-Olig2 (Supplementary Table 1), followed by three washes in 0.1%TritonX-100/PBS and one hour of incubation with the secondary antibodies Alexa Fluor 488 and Alexa Fluor 568 (1:500, Life Technologies). Nuclei were counterstained with DAPI. The structures of MNs were visualized using a Zeiss LSM Confocal Microscope. The percent of positive cells were quantified using ImageJ software and normalised to number of positive staining for Olig2^+^ or SMI-32^+^ or ChAT^+^ on each image.

### Quantitative RT-PCR

RNA was extracted from iPSMNs using the RNeasy Mini Kit (QIAGEN). cDNA was synthesized using the iScript cDNA Synthesis kit (Bio-Rad) with oligodT primers. Quantitative real time PCRs (qRT-PCRs) were performed on 2 μl cDNA using SYBR Fast mastermix (Applied Biosystems) and specific primers for overall *C9orf72* mRNA levels, V1 transcript, V2+V3 transcripts and V3 transcripts, in a total volume of 25 μl per each reaction (Supplementary Table 3). Master mixes and cDNA samples were loaded in a 96-well plates and samples were run on Applied Biosystems StepOne machine and analysed using relative quantitation methods. All samples were amplified in triplicates from at least three different differentiation experiments. All gene expression analysis was performed using ΔΔCt method with normalisation to GAPDH reference gene. Melting curve analysis was used to verify the specificity of the primers. PCR cycling conditions: 95°C 20 seconds initial denaturation, followed by 40 cycles of [95°C 3 seconds, 60°C 30 seconds].

### Promoter methylation assay

For quantitative assessment of methylation levels, quantitative PCR with methylation-sensitive restriction enzymes was applied to measure the C9*orf*72 promoter methylation following published methods [25,26]. Briefly, 100 ng of genomic DNA was digested with 10 units of HhaI (NEB) and 2 units of *Hae*III (NEB) or 10 units of *Hpa*II (NEB) and 2 units of *Hae*III (NEB), to assess the methylation status of two CpG sites in the promoter region. DNA samples digested with only 2 units of *Hae*III were used as a mock reaction. The digestion mixture was incubated overnight at 37°C followed by heat inactivation at 80°C for 20 minutes. qPCR was carried out using 30 ng of digested DNA per reaction and 2X FastStart SYBR Green Master mix (Applied Biosystems) using primers amplifying the differentially methylated *C9orf72* promoter region (Supplementary Table 3). The difference in the number of cycles to threshold amplification between double (*Hae*III and *Hha*I) versus single digested DNA (HaeIII) was used as measure of CpG methylation. Both *Hha*I and *Hpa*II restriction enzymes cut sites are within the unmethylated *C9orf72* promoter and the amplification of the unmethylated digested products after enzyme digestion represents the methylation level.

### Bisulfite sequencing

Direct bisulfite sequencing was achieved using an assay adapted from one described by Xi et al [27]. 400 ng of genomic DNA was bisulfite-converted using the EpiTect bisulfite-converted using the EpiTect bisulfite kit (QIAGEN). PCR was performed on bisulfite-treated DNA using the following primers (as used by Liu et al)[26] to amplify the 5’ CpG island of C9orf72: forward 5’-GGAGTTGTTTTTTATTAGGGTTTGTAGT-3’, reverse 5’-TAAACCCACACCTACTCTTACTAAA-3’. GoTaq Hot Start polymerase (Promega) was used under the following thermal cycling conditions: 95°C for 5 minutes, followed by 10 cycles of touchdown PCR with 30 seconds denaturation at 95°C, 30 seconds annealing reducing the annealing temperature from 70°C to 60°C by -1°C per cycle and with 1 minute extension at 72°C, followed by 30 cycles with the annealing temperature at 60°C and with a final extension step at 72°C for 10 minutes. PCR products of 466 bp were purified by agarose gel extraction and commercially Sanger sequenced by Source BioScience using the forward primer. Resulting chromatograms were analysed using Chromas v2.6 software and the percentage of methylcytosine (mC) determined for each CpG using relative peak heights whereby %mC = C/(C + T). In all samples over 95% of unmethylated C nucleotides were converted to T.

### RNA fluorescence in situ hybridization (FISH)

RNA fluorescence *in situ* hybridisation (FISH) was performed using modified methods previously published [28]. Cells grown on 12mm RNase-free coverslips in 24 well plates were fixed using 4% paraformaldehyde for 20 minutes, washed in PBS and permeabilised using 0.2% Triton X-100 in PBS for 30 minutes. Cells were then dehydrated in a graded series of alcohols, air dried and rehydrated in PBS, briefly washed in 2xSSC and incubated for 30 minutes in pre-hybridisation solution containing 2X SSC and 50% formamide. Hybridisation was carried out for two hours in 250µl of pre-heated hybridisation solution (50% formamide, 2X SSC, 0.16% BSA, 0.8 mg/ml salmon sperm, 0.8 mg/ml tRNA, 8% dextran sulfate, 1.6 mM vanadyl ribonucleoside, 5mM EDTA, 0.2 ug/ml probe) at 80°C. Coverslips were firstly washed three times for 30 minutes each at 80°C in 1ml high-stringency wash solution (50% formamide/0.5X SCC), and then washed three times for 10 minutes each at room temperature in 1ml of 0.5X SCC. This was followed by a brief wash in PBS and immunofluorescence staining with SMI-32 (1:1000, DSHB) was performed to assess the presence of foci in SMI-32+ cells as described above. Optional treatments using RNase A (0.1 mg/ml; Life Technologies: 12091-021) or DNase (150 U/ml, Quiagen: 79254) were performed at 37°C for 1 hour following permeabilisation. After treatment the cells were dehydrated and air dried and the hybridisation protocol was resumed as described above. 2’*O*-methyl RNA sense probes were conjugated with Cy3, whereas antisense probes were conjugated with Alexa Fluor 488 (Integrated DNA Technologies). Probe sequences were (C4G2)_4_, (G4C2)_4_ or (CAGG)_6_.

### RNA Sequencing

RNA was extracted from whole iPSMN cultures at day 30. *Sequin* spike-ins were added to the RNA prior to submission to the sequencing centre, with Mix A added to CRISPR samples and Mix B to *C9orf72* HRE positive iPSMNs. Library preparation was carried out using the Illumina TruSeq RNA Sample Prep Kit v2 by the Oxford Centre for Human Genomics. RNA sequencing was performed on the Illumina HiSeq4000 platform over 75 cycles. Sequences were paired-end. *Fastq* files were generated from the sequencing platform using the manufacturer’s proprietary software. RNA sequencing data from this study have been deposited with NCBI GEO.

### Mapping and Quality Control

Reads were mapped to the custom genome using *STAR* (2.4.2a, [29]) using the two-pass method. Maximum intron size was specified as 2,000,000. During the first run *STAR* was run with an index and junctions generated using the custom genome annotation, as well as the following options: --outFilterType BySJout --outFilterMismatchNmax 999 --outFilterMismatchNoverLmax 0.04 --outFilterIntronMotifs RemoveNoncanonicalUnannotated. The junction file generated in the first run was used in the second run, without additional options. The mapping files from different sequencing lanes were merged at this stage. Quality control was performed on individual *Fastq* files as part of the *cgatflow* pipelines *readqc, rnaseqqc* and *bamstats* [30]. In summary, *fastq* quality was evaluated using *FastQC* (0.11.2) and contamination by other organisms was excluded using *FastQ Screen* (0.4.4). Files were not trimmed. Mapping quality control included *Picard* (1.106) metrics, and plotting of alignment statistics, mapping statistics, library complexity, splicing statistics as well as gene profile plots for assessment of coverage and fragmentation biases.

### Differential expression analysis

Transcripts per million (TPM) and counts tables were obtained using *Salmon* (0.8.2 [31]) with kmer size option 31 and index option fmd. Salmon results were imported into *R* using *tximport* (1.10.0 [32]). Following variance-stabilising transformation of the counts table, exploratory data analysis was performed using heat maps, hierarchical clustering and principal component analysis. Differential expression analysis was performed *DESeq2* (1.22.1 [33]), including the operator in the model, to account for variability across differentiations. Multiple testing correction was performed using the Benjamini-Hochberg correction. The significance of the differential expression results was ascertained using 1000 simulations using random permutations of the group label across all samples with replacement.

### Overrepresentation and enrichment analysis

The results were interpreted using annotations from the gene ontology (GO) database, Kyoto Encyclopaedia of Genes and Genomes (KEGG) database and the mouse genome informatics (MGI) database. RNA Sequins were analysed by providing TPMs and differential expression results tables to *RAnaquin* (1.2.0 [34]). Geneset Enrichment Analysis (GSEA) and was performed using *fgsea* (1.8.0 [35]), and leading edge analysis was performed using the GSEA desktop application [36].

### qPCR for validation

Validation of RNA sequencing results was performed on an extended sample cohort that included 10 genome edited samples, 10 control samples, six C9-02-02 samples from independent differentiations and six patient samples from three different clones from three different patients. Following RNA extraction with the RNeasy Mini Kit (QIAGEN), cDNA was prepared using the High-Capacity cDNA Reverse Transcription Kit (Thermo Fisher Scientific). Primers for qPCR were validated using control iPSC-derived motor neuronal cDNA with 5 serial 1:5 dilutions and included an RT-control. Primer sequences are shown in Supplementary Table 3. qPCR was performed using the Fast SYBR Green Master Mix (Thermo Fisher Scientific) using 0.2μl of cDNA and 250μM primers. 20μl samples were run in triplicate on the Lightcycler480 (Roche) using standard cycling conditions. Three reference genes were used for all reactions (TOP1, RPL13A, ATP5B) and the geometric mean of all three genes was used as the reference for gene expression estimation. Relative abundance estimation was carried out using the 2-ΔΔCt method.

### Western blotting

Protein was extracted for iPSMN cultures at day 30 using RIPA buffer with protease inhibitors and combined with Sample Reducing Agent and LDS sample buffer (Thermo Fisher Scientific). Samples were not heated and run in precast NuPAGE 4-12% Bis-Tris gels (Thermo Fisher Scientific) at 100V for 100 minutes. Protein was transferred on nitrocellulose membranes using the iBlot2 dry blotting system (Thermo Fisher Scientific) and blocked in TBS with 10% skimmed milk containing 0.1% Tween-20. Primary antibodies SV2A (1:1000, Synaptic Systems 119 002) and Syt11 (1:1000, Abcam ab204589) were incubated for 2 hours followed by 3 washes and fluorescent secondary antibody incubation (Licor 800CW and 680RD, 1:10,000). Membranes were visualised in a ChemiDoc imaging system (BioRad).

### Data analysis and statistics for molecular biology experiments

Statistical analyses and data visualisations other than for sequencing analysis were performed using GraphPad Prism.

## Results

### CRISPR/Cas9-mediated gene editing of the *C9orf72* G4C2 repeat expansion

Skin fibroblasts obtained from an ALS/FTD patient (C9-02) carrying an expansion of 1000 G4C2 hexanucleotide repeats in *C9orf72* were used to generate two iPSC lines (C9-02-02 & C9-02-03) as previously described [10]. To correct the HRE mutation and generate isogenic lines, double-nicking CRISPR/Cas9 and HDR was applied to the C9-02-02 iPSC line using a double-stranded DNA (dsDNA) plasmid donor template harbouring two G4C2 repeats and containing two long homology arms flanking each side of the target site, to replace the large repeat expansion (∼1000 repeats) via homologous recombination. A puromycin/TK selection-counterselection cassette was inserted ∼53bp upstream of the repeat region, driven by an EF1a promoter and flanked by two LoxP sites, to facilitate screening and isolation of the corrected clones (Fig. 1A). The cleavage efficiency of the two guide RNAs was evaluated using a T7E1 cleavage assay after transfecting HEK293T cells with either individual gRNAs or with both gRNAs at the same time. The cleavage assay allowed the detection of double-strand breaks, indicated by the presence of heteroduplex DNA (Suppl. Fig. 1A). This demonstrates that both gRNAs were able to introduce DSBs when applied simultaneously, ensuring that they could be used in the subsequent editing experiments in iPSCs.

**Fig. 1.**
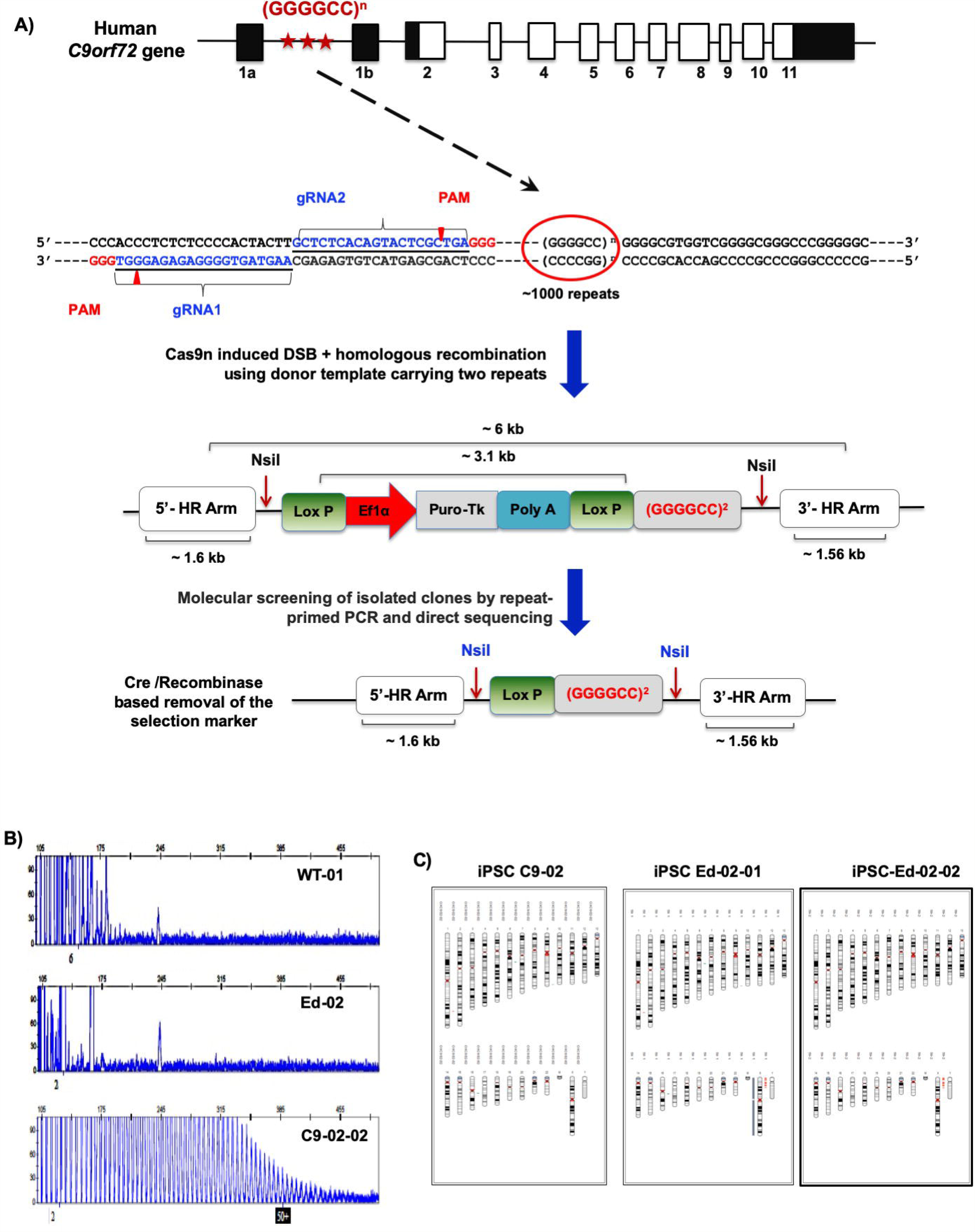
CRISPR Gene targeting strategy for complete correction of the G4C2 repeat expansions in ALS/FTD patient iPSCs. **A.** The overall scheme of the experiment is shown. In the *C9orf72* gene, black boxes represent untranslated exons and white boxes represent translated exons. The position of the G4C2 hexanucleotide repeat region is indicated by stars. The second panel indicates the sequences of both guide RNAs, which are located approximately 140 bp upstream of the repeat region. Double nicking-CRISPR/Cas9 editing was performed on C9-02-02 ALS/FTD patient iPSCs, to remove a repeat expansion (>1000 repeats) and replace it with the wild-type sequence (2 repeats). The homologous donor template design with LoxP-flanked Puro-TK selection cassette for the introduction of normal repeat size is shown in the next diagram, which consisted of ∼1.6kb right homology arm and ∼1.56kb left homogy arm. The selection cassette was inserted directly ∼30bp upstream the repeat region. Cre/LoxP mediated excision was used to remove the selection cassette leaving one copy of LoxP integrated in the genome as demonstrated in the final diagram. **B.** Electropherogram for a healthy control (WT-01), an edited iPSC clone (Ed-02), and the parental C9-02-02 iPSC clone. All Puromycin resistant clones were assessed by repeat-primed PCR (RP-PCR) to identify the presence or absence of the repeat expansion. Twenty-four clones showed normal electropherogram profile. **C.** Karyotyping analysis of the ED-02-01 and Ed-02-02 edited clones revealed a normal chromosome profile compared to the original fibroblast karyotyping result.

Following nucleofection of the C9-02-02 line, puromycin selection was performed and around 100 colonies were isolated and evaluated by repeat-primed PCR to confirm the elimination of the expanded hexanucleotide in 24 clones as indicated by the normal electropherogram profile (Fig. 1B). Sanger sequencing of the repeat region confirmed correct replacement by the donor template in two heterozygous (2% mono-allelic) and three homozygous clones (3% bi-alleic) (Suppl. Fig. 1B). The specific integration of the donor template at the 5’ and 3’ junctions was also evaluated by direct sequencing and restriction fragment length polymorphism (RFLP), confirming specific integration of the donor template (Suppl. Fig. 1C). Bi-allelic clones were excluded due to the manipulation of both alleles with the donor template.

Cre/LoxP mediated excision and negative selection with ganciclovir yielded multiple isogenic lines free of the selection marker, as confirmed by PCR using primers inside the cassette and outside the 5’ homology arm (Suppl. Fig. 1D). After that, two sub clones, Ed-02-01 and Ed-02-02 were selected and the repeat region was amplified and sequenced, confirming the absence of the repeat and presence of the remaining LoxP site in the targeted alleles (Suppl. Fig. 1E&F). The absence of the 6kb expansion in the edited isogenic lines was further confirmed using Southern blotting analysis (Suppl. Fig. 1G). These sequencing results confirmed the presence of the normal repeat size derived from HDR and demonstrating successful editing of HRE mutation by CRISPR/Cas9 system in iPSCs.

Immunostaining of pluripotency markers (NANOG, OCT4 and TRA-1-60) after gene editing showed that Ed-02-01 and Ed-02-02 iPSCs retain normal expression of these markers (Suppl. Fig. 1H). Karyotyping of Ed-02-01 and Ed-02-02 also revealed that both lines retained their normal chromosomal profile when compared to the parental iPSCs (C9-02) (Fig 1C). Immunostaining for motor neuron specific differentiation markers Olig2 and Islet1 on Day 20, and Islet-1, HB9, SMI-32, and ChAT at day 30 showed that the gene editing process had no effect on motor neuron differentiation (Fig. 2A&B), with similar morphology and differentiation efficiencies seen in all differentiated lines (Fig. 2C-F).

**Fig. 2.**
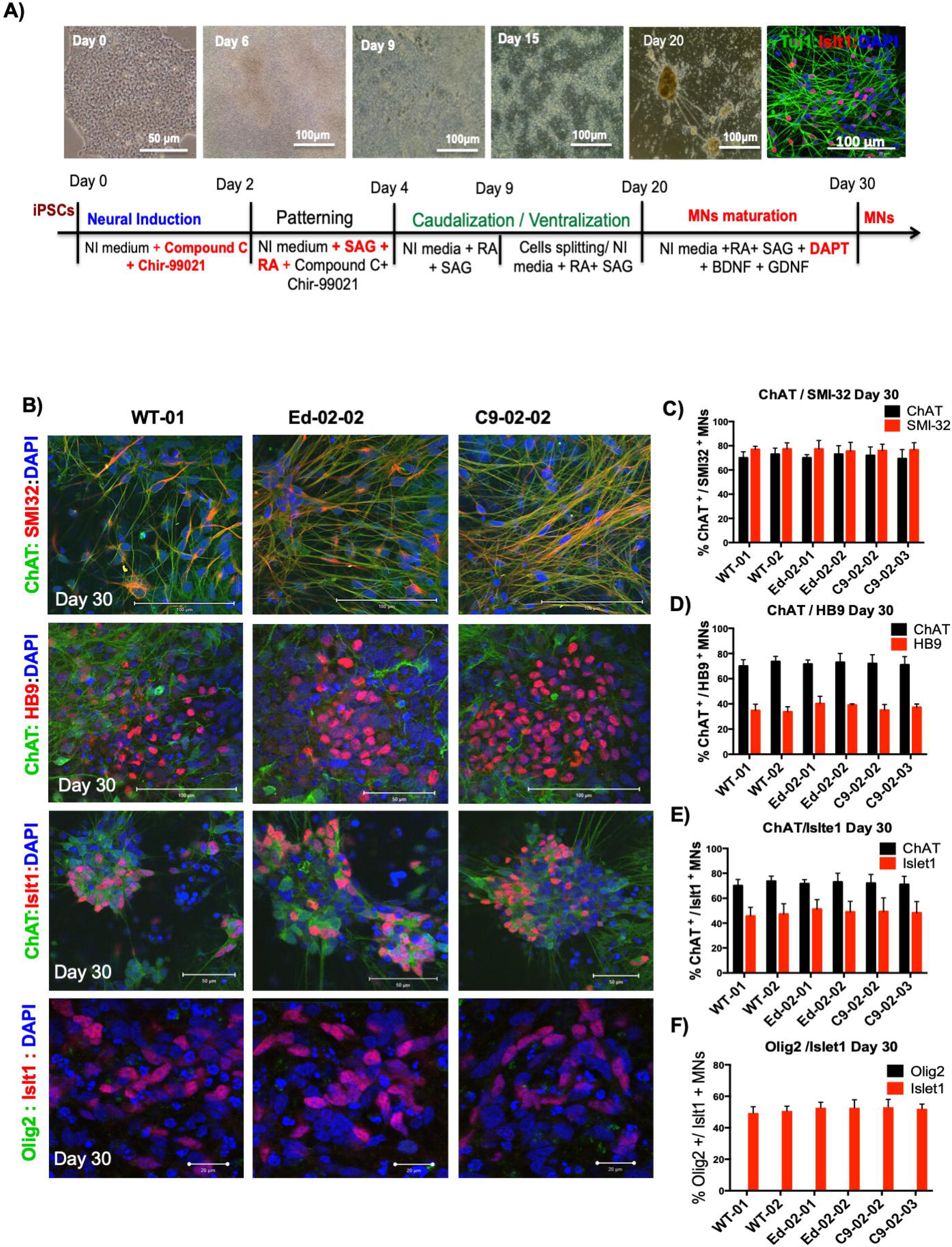
Characterization and motor neuron differentiation of edited clones. **A.** Schematic of the timeline of the MN differentiation protocol and morphology of the cells during the differentiation process. **B.** Representative immunofluorescence images of the MN differentiation markers (Olig2, Isl1, ChAT, SMI32, HB9) derived from WT-01 (healthy control), the Ed-02-02 (edited clone) and the C9-02-02 (*C9orf72* HRE positive clone). **C-F.** Quantification of positive MN specific markers in iPSC-derived MN cultures from two healthy lines (WT-01 and WT-02), two edited lines (Ed-02-01 and Ed-02-02) and two patient lines (C9-02-02 and C9-02-03). On Day 20, 55-60% of the differentiated MNs stained positive for Olig2 and 30-35% of cells stained positive for Islet1, with no difference between the analysed lines (*p* > 0.05). On day 30, 60-70% of MNs population showed positive staining for ChAT, 75-80% of cells stained for SMI32 and 35-40% for HB9 with no differences between the analysed samples (*p* > 0.05, One-Way ANOVA). Olig2 was completely absent on day 30 of differentiation, while Islet1 showed 35-40% nuclear staining. Bar graphs showing mean ± SD. Data from three independent differentiations, minimum of 100 cells per differentiation for each genotype.

### CRISPR/Cas9 genome editing restores normal *C9orf72* expression and DNA promoter methylation levels

Since the HRE in *C9orf72* has previously been shown to reduce the expression of *C9orf72* in iPSC-derived motor neurons, we sought to establish whether normal regulation of expression had been restored by CRISPR/Cas9 genome editing. Quantitative RT-PCR with primers amplifying all *C9orf72* RNA isoforms and the individual variants revealed that total *C9orf72* RNA levels and the levels of the V1 and V2 mRNA variants were reduced in C9-02-02 and C9-02-03 compared to the normalised levels in Ed-02-01 and Ed-02-02, which showed similar levels to WT-01 and WT-02 (*p* < 0.001, One-Way ANOVA) (Fig. 3A).

**Fig. 3.**
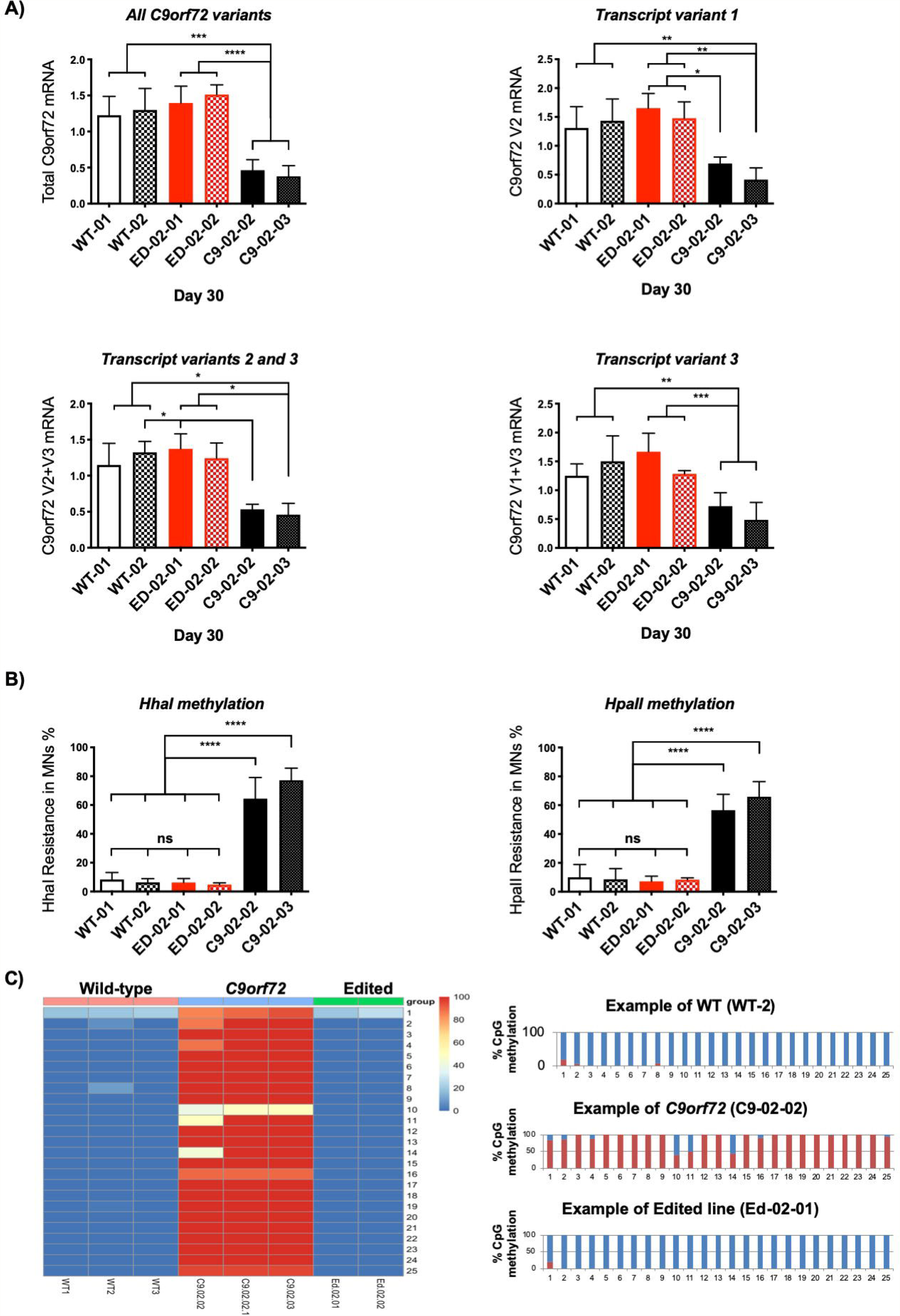
Evaluation of *C9orf72* promoter methylation and gene expression. **A.** A reduction in Total, V1 and V2 *C9orf72* RNA is seen in C9-02-02 and C9-02-03 clones compared to healthy controls and Edited lines Ed-02-01 and Ed-02-02 (*** *p* < 0.001, ** *p* < 0.01, Bonferroni’s multiple comparison’s test). The results suggest restoration of normal transcription at the *C9orf72* locus following genome editing. Data showing mean ± SD from three independent differentiations. **B.** Following digestion with the methylation sensitive restriction enzymes Hha1 and HpaII, CpG island hypermethylation is demonstrated in the *C9orf72* promoter region of patient lines C9-02-02 and C9-02-03 compared with both controls WT-01 and WT-02 and edited lines Ed-02-01 and Ed-02-02 (**** *p* < 0.0001, Bonferroni’s multiple comparison’s test). Data showing mean ± SD from three independent differentiations. **C.** Bisulfite sequencing of all CpG islands in the 5’ region upstream of the *C9orf72* HRE demonstrated hypermethylation in the *C9orf72* positive lines. In the two edited lines studied, this was restored to normal. The right panel displays three example plots showing the ratios of methylated to unmethylated cytosines at each CpG locus.

Previous studies have also shown that *C9orf72* gene expression may be reduced by repeat-associated methylation of the CpG promoter [37,38]. To evaluate methylation levels upstream of the repeat, we performed a methylation sensitive restriction enzyme-qPCR which has previously been described [25,26]. Restriction digestion of the CpG site by HhaI and HpaII occurs in the absence of CpG DNA methylation. Quantification of the digested DNA derived from both enzymes showed that CpG methylation was higher in C9-02-02 and C9-02-03 patient lines compared to WT-01 and WT-02 healthy controls and Ed-02-01 and Ed-02-02 lines for HhaI (*p*< 0.0001) and HpaII digestion *(p* < 0.0001) (Fig. 3B).

In order to ascertain the methylation status of multiple CpG sites within the 5’ CpG island region of *C9orf72*, genomic DNA samples isolated from iPSMNs clones were treated with sodium bisulfite and subjected to PCR using primers specific to the 5’ CpG region. Direct Sanger sequencing of PCR products allowed relative quantification of C versus T nucleotides at each CpG site as a measure of methylation, since unmethylated C is converted to T by bisulfite, while methylated C (mC) remains unchanged. A total of 26 CpG sites could be assessed by this method, including one site (CpG 5) formed by an A>G SNP positioned 151 nucleotides upstream of the exon 1a start site (rs1373537, hg38 9:27574017T>C), which was present in all DNA samples analysed in this study. Although the C allele of this SNP is not present in the reference sequence, it is in fact the predominant genotype in all populations studied, with the T allele having a minor allele frequency of only 0.01 as determined by the 1000 Genomes Project.

Using this method, all 26 CpG residues from iPSC motor neuron clones derived from *C9orf72* HRE-positive patients were found to be methylated (>50% mC). In contrast, the same CpG residues were found to be unmethylated (< 30% mC) in control iPSMNs clones and in edited clones. Notably, the majority (86%) of CpG residues in the analysed samples were either 100% or 0% methylated. CpG 1 was the site most likely to have a mixed methylation pattern, being mixed in every sample tested. Note that in this assay, CpG 2 corresponds to the site assayed by the methylation-sensitive restriction digestion assay (Fig. 3C).

Collectively these results suggest that gene editing of the repeat expansion resulted in normalisation of *C9orf72* gene expression levels through restoration of normal methylation patterns.

### RNA foci and polydipeptides are abolished in CRISPR/Cas9 corrected iPSMNs

The presence of RNA foci and toxic polydipeptides are established neuropathological hallmarks of the *C9orf72* HRE expansion and can be detected in iPSMNs derived from affected patients. We evaluated the presence of sense and antisense RNA foci in the iPSMNs using fluorescence in situ hybridisation (FISH). Both sense and antisense RNA foci were seen in motor neurons derived from C902-02 and C9-02-03 patient lines but were absent from healthy controls WT-01 and WT-02 and edited clones Ed-02-01 and Ed-02-02 (Fig. 4A-C).

**Fig. 4.**
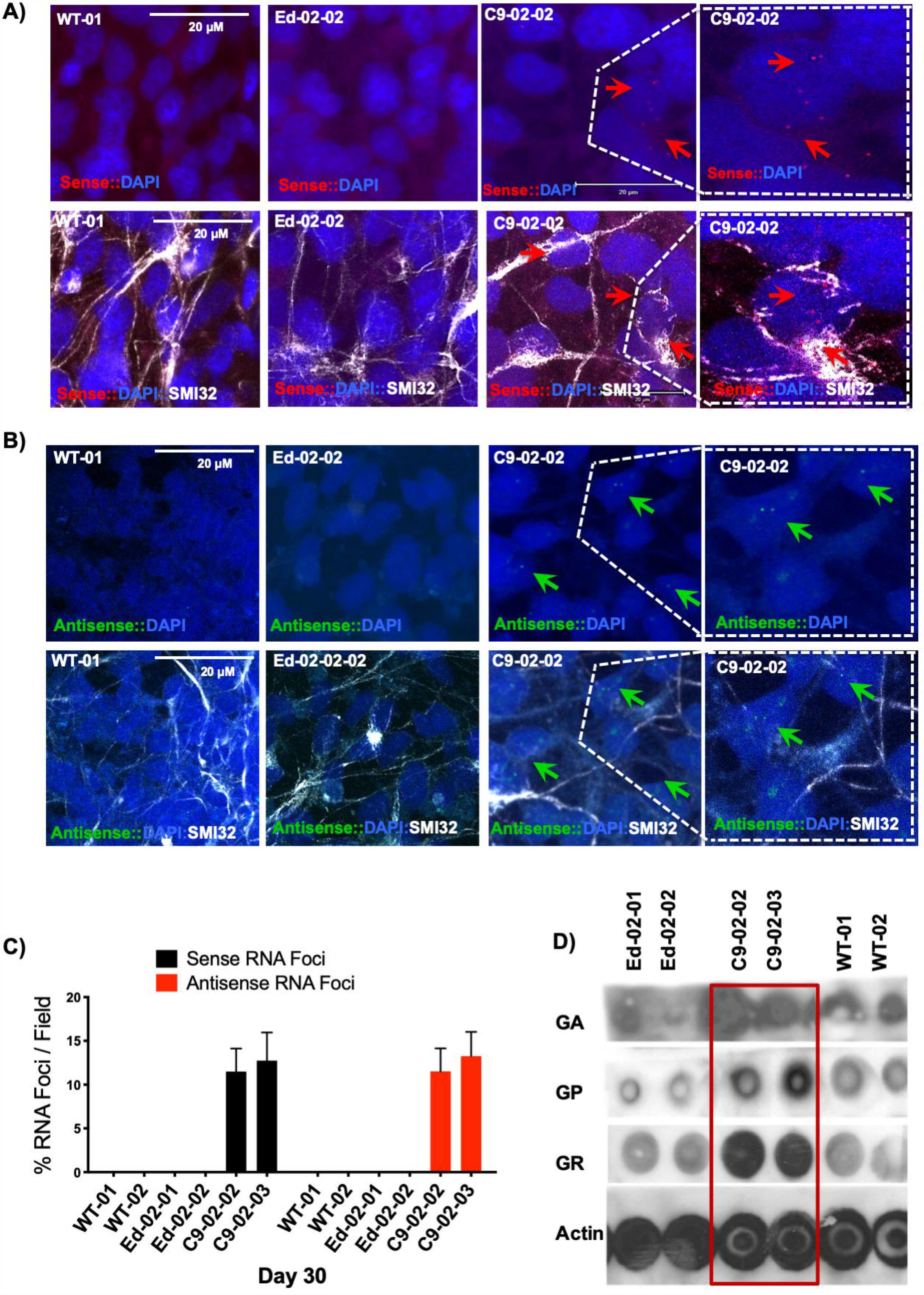
Abolition of sense and antisense RNA foci and reduction of RAN translation products. **A&B.** Fluorescence in situ hybridisation with G_2_C_4_-Cy3 and G_4_C_2_-Alexa488 probes showing G_2_C_4_ sense (red) G_4_C_2_ antisense RNA foci (green) in C9-02-02 iPSC-derived MNs. Foci are absent in WT-02 healthy and Ed-02-02 edited clones. Nuclear foci are indicated by arrows and SMI32 was used as a MN-specific marker. **C.** The percentage of positive nuclei for sense and antisense RNA foci was between 10-12%, with no foci detected in edited and healthy clones. **D.** Dot-blot analysis shows higher concentrations of GA, GP, and GR dipeptide repeats in C9-02-02 and C9-02-03 compared to both healthy controls and edited lines.

To examine whether the edited MNs rescue the formation of RAN-dipeptides (DPs), we performed a dot-blot analysis of the GA, GP, and GR polydipeptides using protein samples from all experimental lines. Higher concentrations of the DPs were detected in iPSMNs of C9-02-02 and C9-02-03 lines compared to healthy and edited lines (Fig. 4D). These results confirm that removal of the G_4_C_2_ HRE successfully corrects the typical pathology of nuclear RNA foci and cytoplasmic dipeptides in iPSMNs.

### CRISPR/Cas9 correction improves cell survival and the response to cellular stress

In previous work we identified increased susceptibility of *C9orf72* HRE positive iPSMNs to apoptosis in basal conditions [10]. In this study we confirmed increased levels of cleaved caspase3 in *C9orf72* HRE positive iPSMNs using immunofluorescence imaging and demonstrate that edited iPSC-derived motor neurons, Ed-02-01 and Ed-02-02, showed a decrease in cleaved caspase-3 staining compared to patient lines C9-02-02 and C9-02-03 (Fig. 5A, *p <* 0.001).

**Fig. 5.**
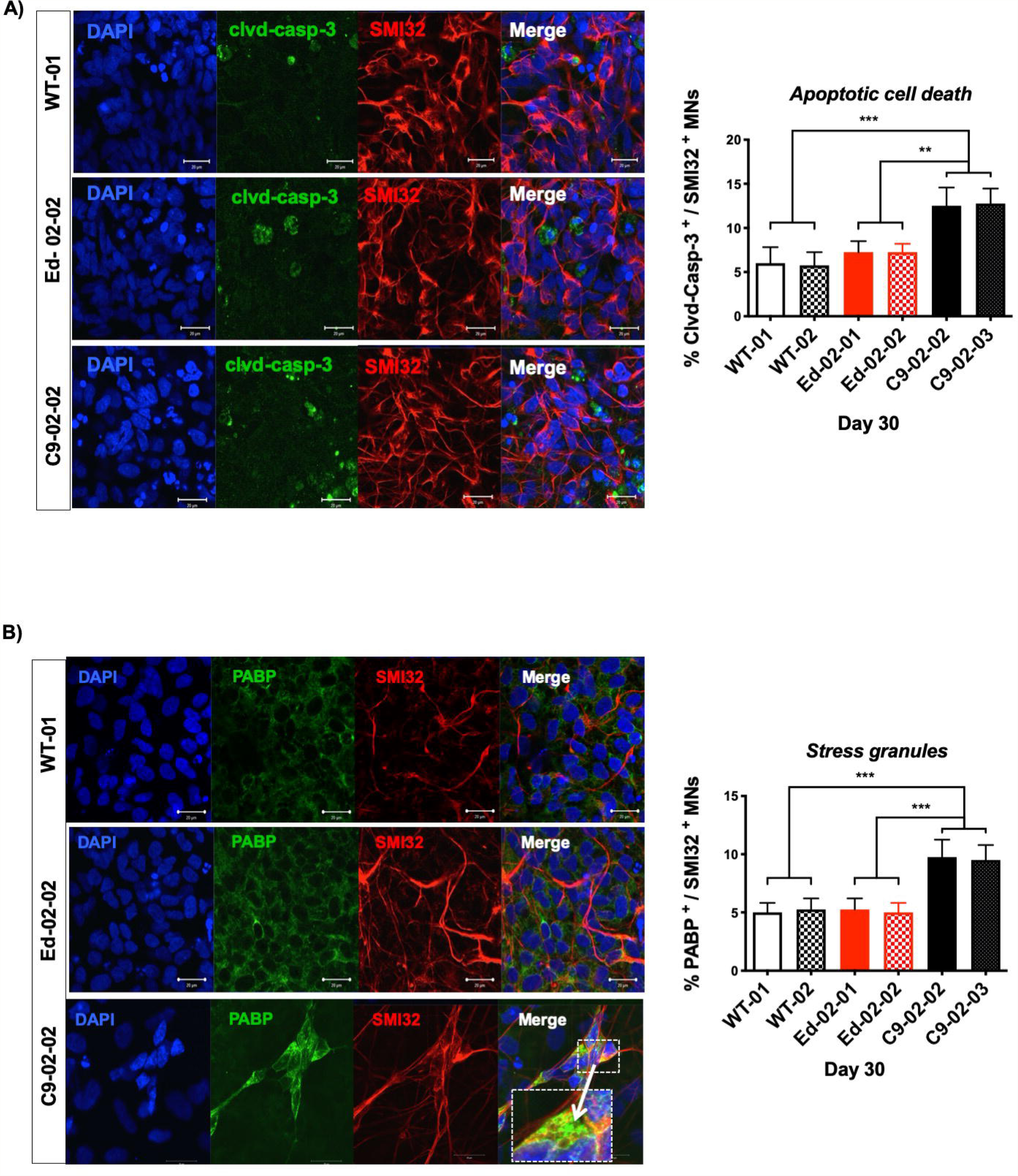
Edited iPSMNs are less susceptible to apoptotic cell death and glutamate excitotoxicity. **A** Representative images of cleaved caspase-3 activation in iPSC-derived motor neurons from lines WT-01, ED-02-02 and C9-02-02 at basal conditions. Quantification of the proportion of cleaved caspase-3 positive motor neurons (mean ± SD from three independent differentiations, minimum 100 cells per line and differentiation) demonstrates a decrease in cleaved caspase-3 frequency in Ed-02-01 and Ed-02-02 cells compared to C9-02-02 and C9-02-03 iPSMN cultures (\*\**p* < 0.01, *** *p* < 0.001, Bonferroni’s multiple comparison’s test). **B.** PolyA binding protein (PABP) accumulation in stress granules was detected at higher frequency in C9-02-02 and C9-02-03 patient iPSC-MNs compared to healthy controls WT-01 and WT-02 as well as edited lines Ed-02-01 and Ed-02-02 clones (* *p <* 0.05, **** *p* < 0.0001, Bonferroni’s multiple comparison’s test, mean ± SD from three independent differentiations, minimum 100 cells per line and differentiation).

We have previously reported that the *C9orf72* iPSC-derived motor neurons show a reproducible increase in markers of cellular stress, including a higher frequency of PABP positive stress granules. Compared to C902-02 and C9-02-03 lines, the number of polyA-binding protein (PABP) positive stress granules in Ed-02-01 and Ed-02-02 iPSC-derived MNs was restored to levels similar to control lines (Fig. 5B, *p* < 0.001).

### RNA sequencing confirms *C9orf72* haploinsufficiency and retention of the repeat-containing intron in *C9orf72* HRE carriers

To enable an unbiased appraisal of the biological consequences of the hexanucleotide expansion, we performed RNA sequencing of iPSMN cultures at 30 days differentiation. We sequenced a total of seven edited samples (4 Ed-02-01 and 3 Ed-02-02), four samples with the hexanucleotide expansion (all C9-02-02) and one control patient sample. All differentiations were performed independently by a total of three operators. GC content was 43.4 % (SD 0.7), 5’ to 3’ bias obtained from Picard tools was 0.36 (SD 0.07) and read depth was 56.6 million (SD 4.6 million), and these did not differ between groups (*p* > 0.05). Further quality control was performed by the addition of “sequin” spike-ins that showed a good correlation between actual and predicted concentrations (Suppl. Fig. 2A).

On exploratory data analysis, there was separation between C9-02-02 and edited controls in principal component 1, after correcting for operator batch effect in the model (Fig. 6A). Motor neurons from this differentiation had a rostral identity, corresponding to cervical HOX gene expression and in line with other studies (Suppl. Fig. 2B). Pluripotency markers were not expressed (Suppl. Fig. 2C) and motor neuron markers were expressed in all cell lines (Suppl. Fig 2D).

**Fig.6.**
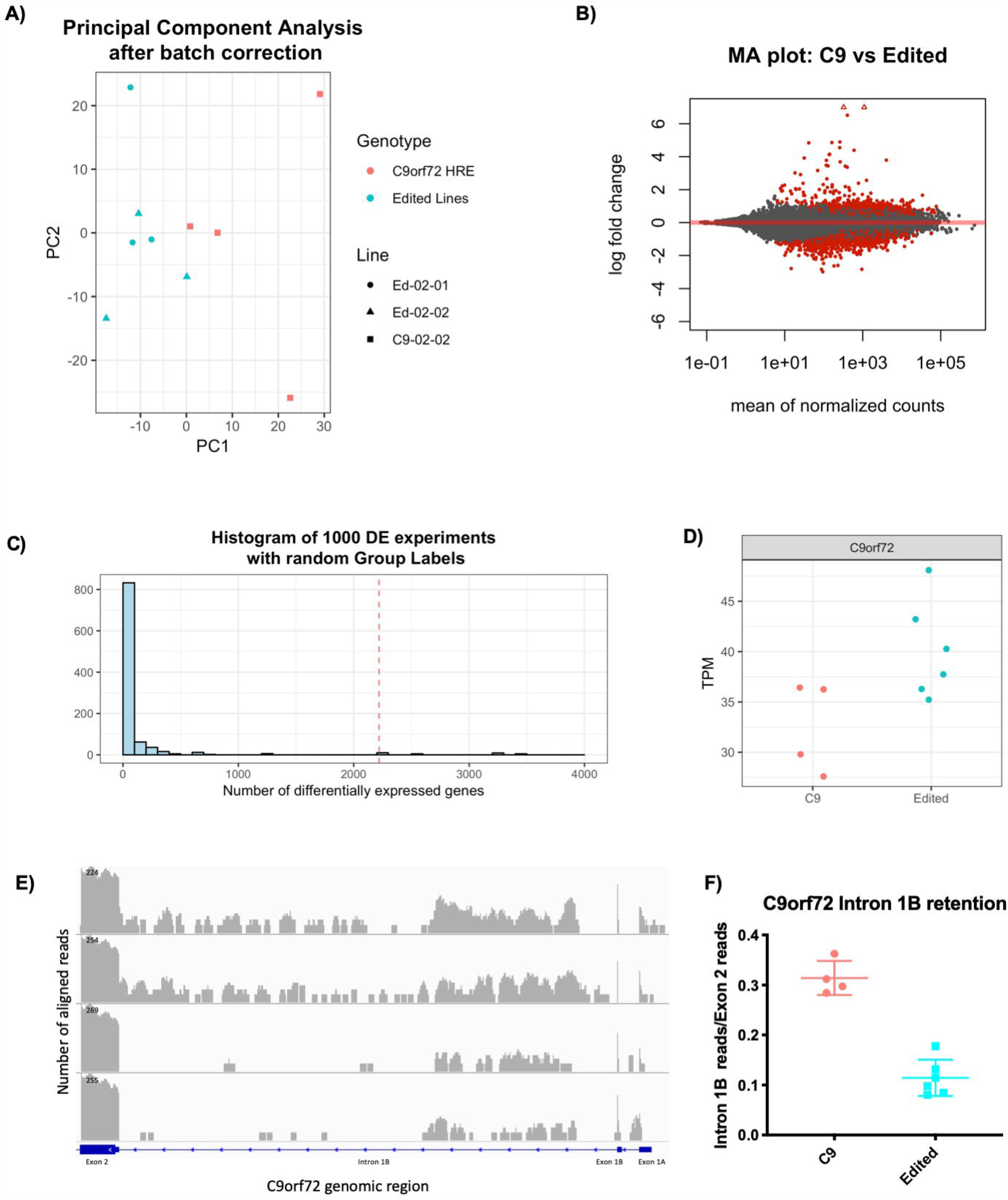
Genome editing of *C9orf72* results in significant differential expression and reduced retention of the repeat containing *C9orf72* intron. **A.** Principal component analysis using the 500 genes with the largest variance shows separation between the *C9orf72* HRE positive parent line and both edited controls. **B.** Differential expression analysis comparing the C9-02-02 clone with both edited clones reveals 1013 upregulated and 1203 downregulated genes as seen on this MA plot. **C.** Histogram of permutation analysis using shuffled line labels demonstrates statistical significance of the sequencing analysis (p = 0.034). Histogram indicates frequency of having N differentially expressed genes upon permutation. Dashed red line indicates result without permutation. **D.** Transcripts per million (TPM) showing normalised expression for all *C9orf72* transcripts combined (FDR = 0.002). **E.** Representative images showing stacked reads aligned to the first three exons and corresponding introns of *C9orf72* (log scale). Top two panes show two independent differentiations of line C9-02-02 and the lower two panes show reads from Ed-02-01 and Ed-02-02. The gene diagram is annotated with the used nomenclature. **F.** Quantification of intron retention of the repeat containing intron divided by the number of reads in the adjacent exon 2 shows reduced intron retention after genome editing (Mean ± SD, *p* = 0.01, Mann-Whitney test).

On gene level differential expression analysis, 2220 genes were found differentially expressed at an FDR threshold of 0.05, of these 1017 were upregulated and 1203 were downregulated (Fig. 6B). Permutation analysis of differential expression was statistically significant (*p* = 0.027, Fig. 6C). Total *C9orf72* expression was reduced by 22.5% in *C9orf72* HRE motor neurons compared to the edited controls, confirming previous qPCR findings (FDR = 0.002, Fig 6D). Finally, in line with other studies [39], we detected retention of the repeat containing intron in the *C9orf72* HRE positive lines which was restored to lower levels in the edited lines (*p* = 0.01, Mann-Whitney test, Fig 6 E,F). On analysis of splicing changes throughout the genome using *rMATS* (version 4.0.1) no statistically significant differential alternative splicing events were found (*p* = 0.2).

### Enrichment of ALS-relevant pathways including synaptic function in *C9orf72* HRE positive motor neurons

Overrepresentation analysis revealed striking differences in up- and downregulated genes. Genes upregulated in *C9orf72* HRE positive motor neurons were enriched for calcium-ion dependent exocytosis, synapse organization and neurotransmitter transport, whereas downregulated genes were enriched for cell division (Fig 7A).

**Fig. 7.**
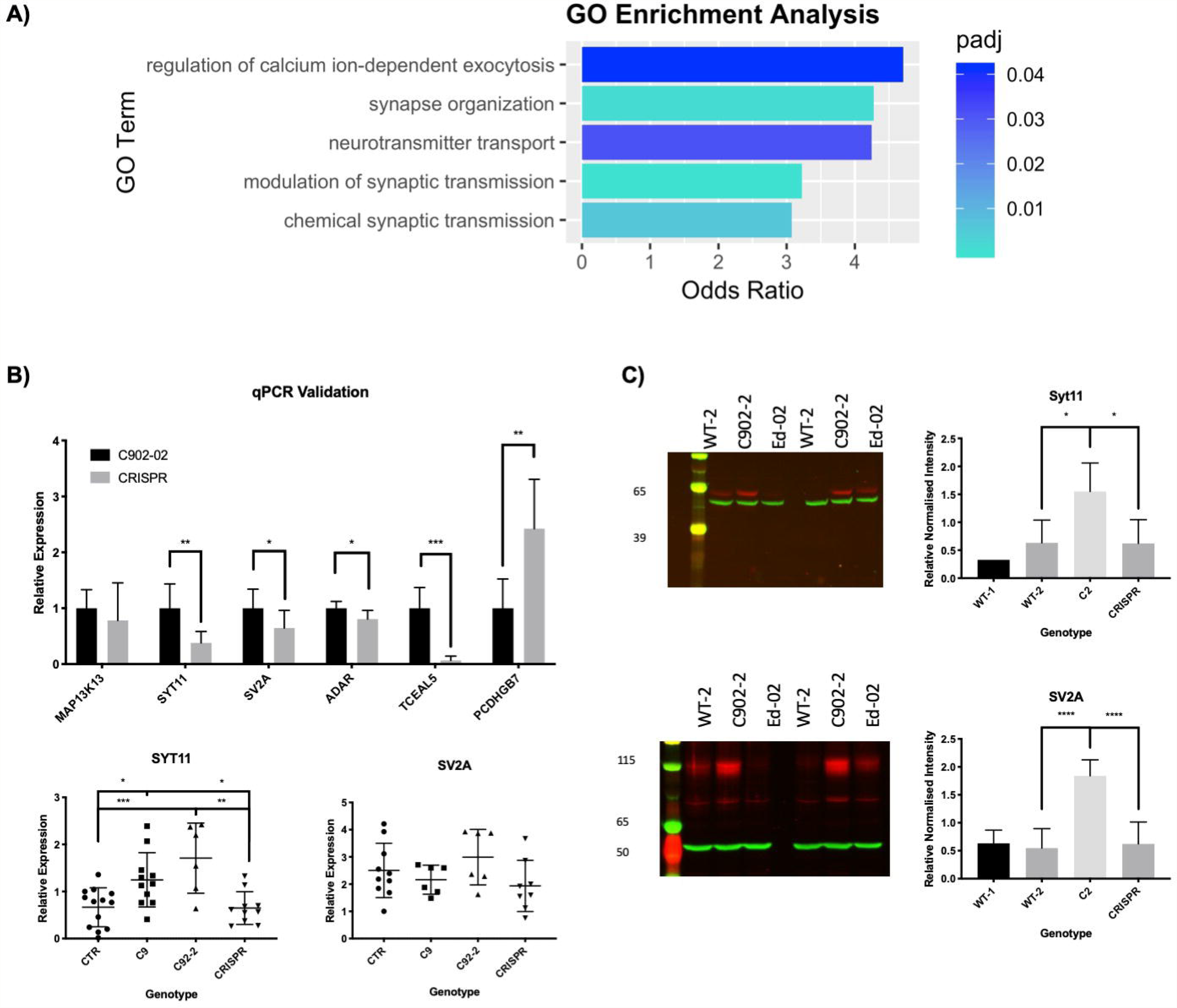
Geneset and overrepresentation analyses of gene expression increased in *C9orf72* HRE neurons compared to edited controls reveals pathways relevant to ALS. **A.** Gene ontology overrepresentation analysis of genes upregulated in *C9orf72* HRE iPSMNs compared to the edited controls (FDR < 0.05) reveals enrichment of pathways of calcium-ion dependent exocytosis and synaptic transmission**. B.** Geneset enrichment analysis (GSEA) using the reactome datasets indicates enrichment of glutamate binding and relevant signal transduction pathways in *C9orf72* HRE positive neurons, independently of the p value threshold. **C.** GSEA using curated genesets from the MSigDB C2 set reveals enrichment of the KEGG Amyotrophic Lateral Sclerosis Dataset driven largely by the relatively increased expression of glutamate-receptors, neurofilaments as well as BAX and Bcl-XL. **D.** Leading edge analysis of all enriched reactome pathways demonstrates overlap between the various pathways, suggesting the presence of a core regulon driving these pathways. **E.** A number of differentially expressed targets were selected for validation of RNA sequencing. Validation was performed on six C9-02-02 lines and seven edited lines (Mean ± SD; Mann-Whitney test: \*\*\**p* < 0.001, ** *p* < 0.01, * *p* < 0.05). For genes of interest, more extensive validation was undertaken as follows: CTR: WT-1 5 differentiations of one line, and WT-2 8 differentiations of one line; C9: 2 differentiations of C9-01, 4 differentiations of C9-02-03, 3 differentiations of C9-04 and 2 differentiations of C9-0T; C9-02-02: 6 differerentiations; CRISPR: 5 differentiations of Ed-02-01 and 4 differentiations Ed-02-02. (Mean ± SD; Bonferroni’s post-hoc test: * *p* < 0.05, *** *p* < 0.001). **F.** Six independent differentiations of control lines (WT-01 and WT-02), edited lines (C9-02-02 and C9-02-03) and control lines (WT-01 and WT-02) were performed for western blotting using Synaptotagmin 11 and SV2A antibodies. The left panels show quantification across all six differentiations (Mean ± SD). Synaptotagmin 11 has increased expression in C9-02 lines compared to edited lines (*p* = 0.04, Bonferroni’s post-hoc test) and healthy controls (*p* = 0.04, Bonferroni’s post-hoc test). SV2A is increased in C9-02 lines compared to edited lines (*p* < 0.001, Bonferroni’s post-hoc test) and healthy controls (*p* < 0.001, Bonferroni’s post-hoc test). The right panels show representative western blots for both proteins (actin green, antibody of interest red).

To corroborate these findings using a different method that does not depend on a p-value threshold, geneset enrichment analysis using the reactome database was performed. The top terms enriched in motor neurons from *C9orf72* HRE carriers included Ras activation upon Ca^2+^ influx and neurotransmitter release (Suppl. Fig. 3A). Interestingly, the KEGG Pathway for amyotrophic lateral sclerosis was also enriched in *C9orf72* HRE positive iPSMNs (adjusted p-value 0.03). The pathway contains genes involved in a number of ALS relevant pathogenic processes such as glutamate toxicity, mitochondrial dysfunction, ubiquitin proteasome dysfunction as well as neurofilament damage (Suppl. Fig. 3B). Leading edge analysis using all enriched reactome genesets (adjusted *p* < 0.05) was performed, to look for genes that are enriched in multiple pathways and therefore may be involved in pathway regulation (Suppl. Fig. 3C). This analysis yielded a group of genes shared by four or more pathways which included AMPA and NMDA glutamate receptor subunits, synaptic genes, neurofilament light chain and Ras nucleotide exchange factors (Suppl. Fig. 4A). There was strong overlap between these genes and the leading edge of the ALS KEGG pathway (Suppl. Fig. 4B). Of relevance to functional results indicating caspase3 activation, the KEGG ALS pathway analysis results included an enrichment of ALS-relevant apoptotic pathways including an upregulation of BAX (FDR = 0.04, log2-fold change 0.8).

Validation of the RNA sequencing study by qPCR was undertaken in an expanded sample cohort and confirmed the findings from the sequencing study (Fig. 7B). For further validation, we focussed on the enrichment of transcripts encoding synaptic proteins in *C9orf72* HRE positive motor neurons. The synaptic constituents Synaptotagmin 11 was found to be consistently upregulated in *C9orf72* HRE iPSMNs compared to both isogenic controls and healthy controls. Synaptotagmin 11 transcript levels in other *C9orf72* HRE positive iPSMN lines (combined data from three clones from three different patients) were similarly elevated (Fig 7B).

To study whether the increased levels of Synaptotagmin 11 affect protein expression we performed western blotting on total soluble protein obtained from six independent motor neuron differentiations and were able to confirm increased levels of Synaptotagmin 11 and SV2A in C9-02 iPSMN lines compared to both healthy controls and edited lines (Fig. 7C), demonstrating that gene editing has restored the expression of these proteins to their baseline levels.

## Discussion

Modelling ALS using iPSC-derived motor neurons offers novel opportunities in studying human motor neurons *in vitro* and has substantially contributed to the understanding of *C9orf72* HRE positive ALS [10,15,40]. Key challenges in the interpretation of expression changes in iPSC based models include genetic differences between individuals and also biological variance between cell lines from the same individual, which might mask true expression changes related to the disease mutation. This makes it challenging to relate changes in transcription to cellular phenotypes to individual mutations, and to the underlying pathophysiology of ALS. The complex genetic architecture of ALS, in which even apparently monogenic forms of the disease may have an oligogenic contribution, makes it even more challenging to isolate the specific impact of individual mutations on cellular phenotypes relevant to ALS [41,42]. Genome editing, particularly using the CRISPR/Cas9 nuclease, provides a method to address these challenges directly by creating isogenic control lines, and has been successfully used to correct ALS-causing point mutations [17–19].

The CRISPR/Cas9 system is a very versatile and simple RNA-mediated system for genome editing in diverse cell types and organisms, with the major advantage of the capacity to target virtually any gene in a sequence-dependent manner [43]. Targeting of Cas9 to a specific genomic site is mediated by a 20-nucleotide guide sequence within an associated single guide RNA (sgRNA) which directs the Cas9 to cleave complementary target DNA-sequences adjacent to short sequences known as protospacer-adjacent motifs (PAMs). Cas9 introduces a double-strand break (DSB) 3-base pairs upstream of the PAM, which is repaired by either of two pathways, non-homologous end-joining (NHEJ) which leads to insertion/deletion (indel) mutations of various lengths, or by homology-directed repair (HDR) which can be used to introduce specific mutations or sequences through recombination of the target locus with exogenously supplied DNA ‘donor’ templates [22].

Using this approach, Selvaraj et al. have reported the excision of the *C9orf72* HRE and its flanking region in three iPSC lines from different patients demonstrating the correction of neuropathological features in iPSMNs, as also demonstrated in this study [16]. Among several hundred transcriptional changes, they demonstrate increased GluA1 AMPA receptor expression in patient compared to edited lines. Editing resulted in a reduction of MN-specific vulnerability to excitoxicity. The three lines in the study were corrected using non-homologous end joining, resulting in deletions of variable extent in the promoter region of the most abundant *C9orf72* transcript. A second study using a similar approach in two cell lines, demonstrated upregulation of proapoptotic pathways including ATM and p53 that were corrected by excision of the repeat [44].

Our study uniquely uses homology directed repair to reintroduce a donor template from the other allele, thereby avoiding the deletion of promoter RNA, and leaving behind two short LoxP insertions. We demonstrate that this method results in restoration of normal *C9orf72* expression and correction of repeat-associated hypermethylation of CpG islands, as well as in reduced intron retention. Similarly, we also observe an increase in Ca^2+^ permeable subunits of AMPA receptors in patient cells, suggesting increased glutamate toxicity.

In line with previous results [10,44], this study also demonstrates increased caspase-3 cleavage in *C9orf72* HRE positive iPSMNs, which is restored to normal levels after genetic correction. Cleaved caspase-3 is an indicator of ER stress and apoptosis, and has been associated with gain of toxic function in poly-GA overexpressing mouse cortical neurons [45]. The pro-apoptotic milieu associated with the *C9orf72* HRE is further corroborated by RNA sequencing results, which show increased *BAX* expression in patient iPSMNs and is in line with our previous findings of reduced Bcl-2 and increased levels of BIM and Bcl-X_L_ [10].

In summary, this study provides further evidence for specific pathways underlying the pathogenicity of the *C9orf72* HRE expansion by restoring normal cellular phenotypes following focused correction of the expansion using homology directed repair of the *C9orf72* HRE. The successful correction of both gain and loss of function mechanisms demonstrates complete correction of the consequences of the HRE, which has therapeutic application in the longer term, however the low efficiency of HDR in mature neurons would first need to be overcome. RNA sequencing further confirms the correction of pathways relevant to ALS and offers new potential targets for investigation such as upregulation of synaptic proteins seen in this study.

## Supporting information

Supplementary Figure Legends, Tables and Sequences

## Acknowledgements

We thank the High-Throughput Genomics Group at the Wellcome Trust Centre for Human Genetics (funded by Wellcome Trust grant reference 090532/Z/09/Z) for the generation of the sequencing data.

We would like to thank Dr Timothy R. Mercer and the Garvan Institute of Medical Research for the provision of sequin spike-ins.

## Author Contributions

N.A., J.S., R.F., A.D., D.S, R.D., S.C. and K.T. conceived and planned the experiments. N.A., J.S., R.F., R.D and A.D. carried out the experiments. J.S., M.T. and K.T. provided patient samples. J.S. and N.A. wrote the paper. All authors provided critical feedback on the manuscript.

