## Supplementary figures and images for "Correction of amyotrophic lateral sclerosis related phenotypes in induced pluripotent stem cell-derived motor neurons carrying a hexanucleotide expansion mutation in *C9orf72* by CRISPR/Cas9 genome editing using homology-directed repair"

### Supplementary Figure 1

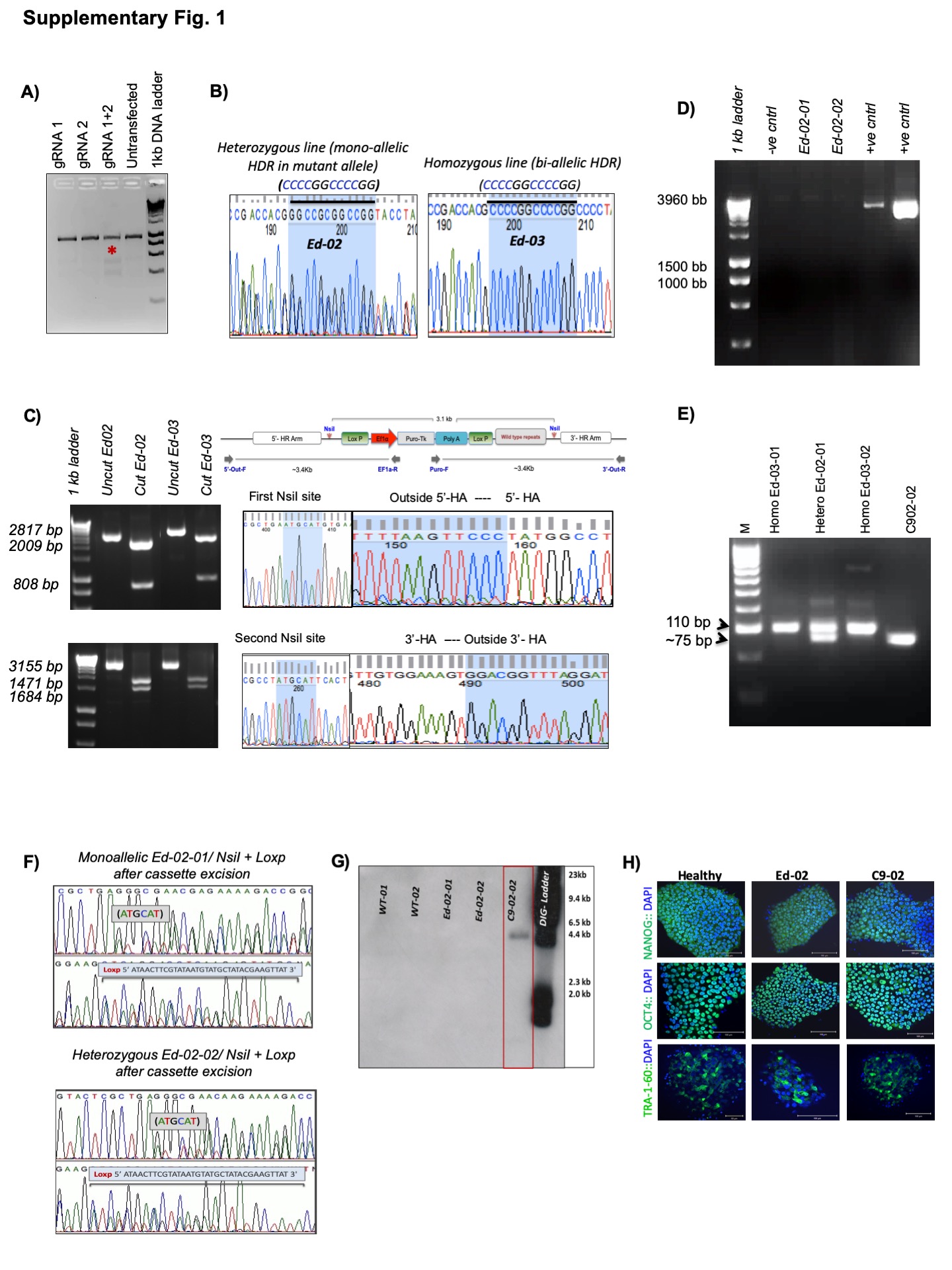

### Supplementary Figure 2

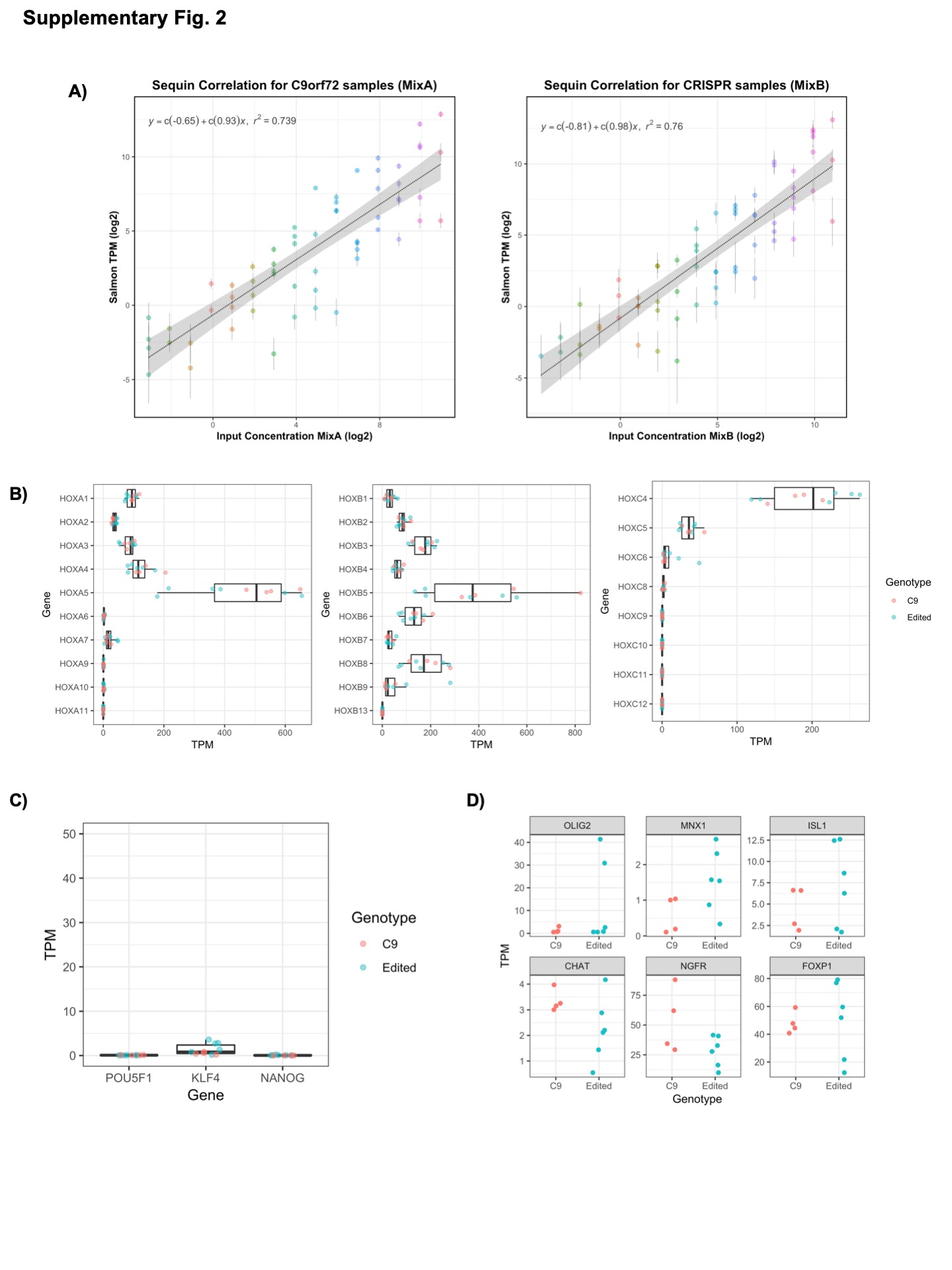

### Supplementary Figure 3

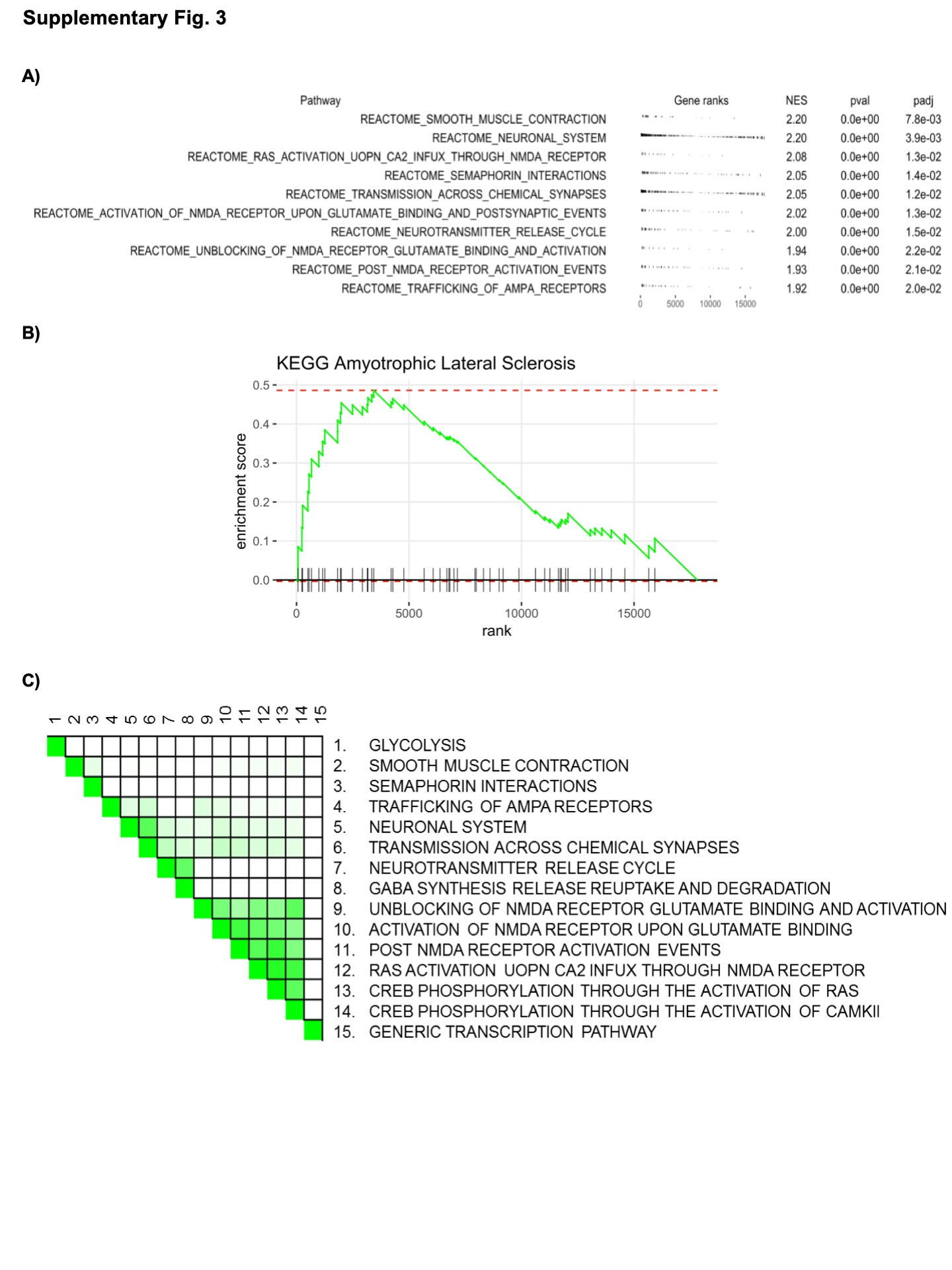

### Supplementary Figure 4

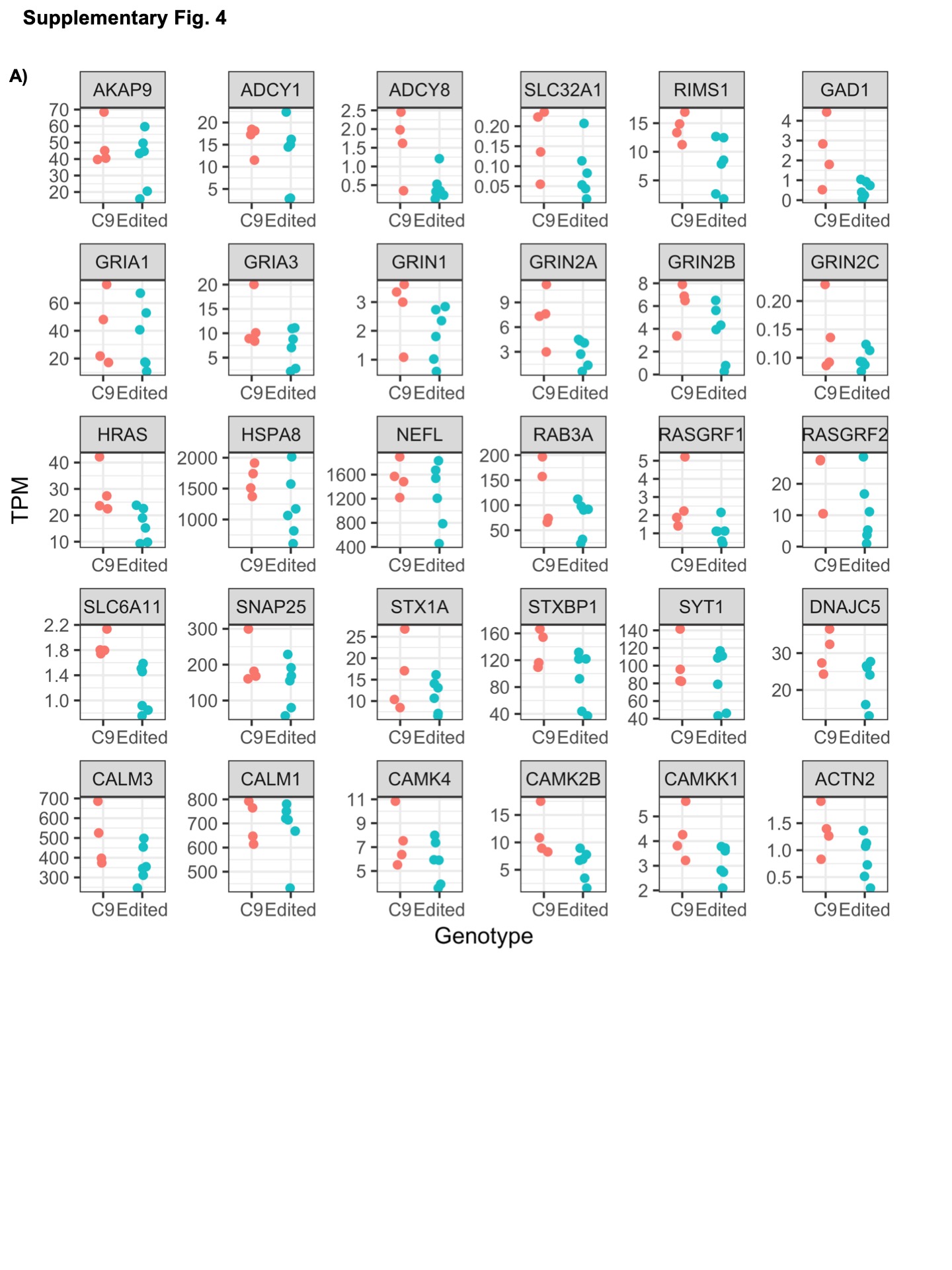

### Supplementary Figure 5

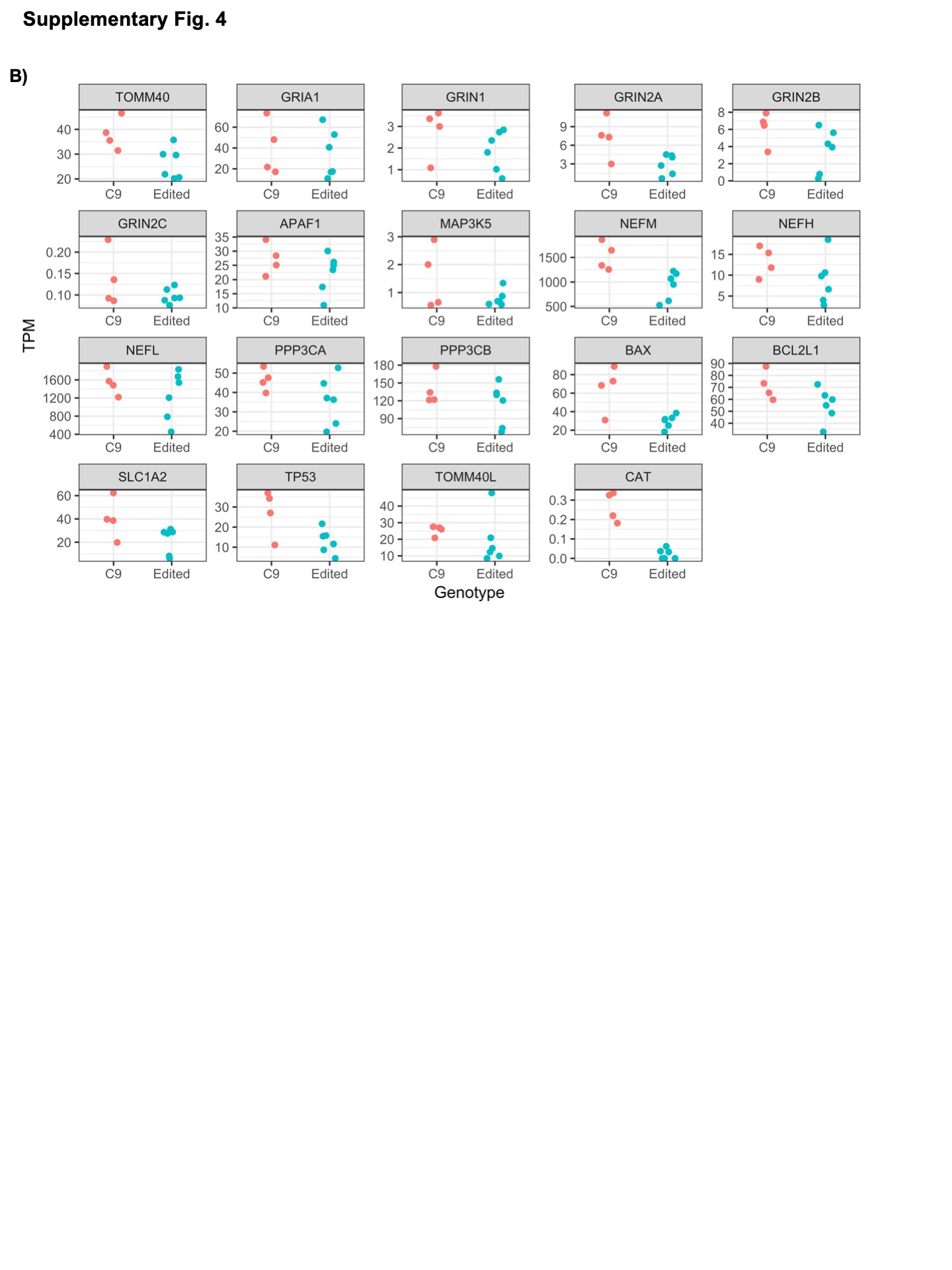
