## Supplementary Figure Legends, Tables and Sequences for "Correction of amyotrophic lateral sclerosis related phenotypes in induced pluripotent stem cell-derived motor neurons carrying a hexanucleotide expansion mutation in *C9orf72* by CRISPR/Cas9 genome editing using homology-directed repair"

**Fig.S1. Detailed molecular evaluation of CRISPR/HDR targeted ED-02 cells and successful removal of the selection cassette.** **A.** T7EI cleavage assay on Hek293T cells was performed, to evaluate the efficiency of the gRNAs in making DSBs indicated by the red star. **B.** Sequencing of the repeat region to confirm the presence wild-type repeat size. The first image shows a mono-allelic targeted clone with HDR only in the mutant allele (Ed-02). The second image shows a bi-allelic targeted clone with HDR in both alleles (Ed-03). **C.** PCR-RFLP and sequencing of the region upstream and downstream the repeat to check for site-specific integration of the donor template into 5’ and 3’ junctions. The primers for PCR are indicated by the arrows (5’out-F, Ef1a-R, Puro-F, and 3’Out-R). **D.** Agarose gel result showing the deletion of the selection marker by a lack of amplification with primers inside the Puro-Tk cassette and outside the 5’ junction, the original clone carrying the puromycin cassette was used as a positive control. **E.** Successful removal of the puromycin cassette by the amplification of the repeat region using primers covering the LoxP site. The wild-type allele results in a 75bp fragment and the targeted allele produces a 109 bp fragment (including the remaining 34bp LoxP site). **F.** Sequencing confirmation for Ed-02-01 & Ed-02-02 after the removal of the selection marker. Sequence analyses revealed correction of the repeat expansion and successful insertion of the wild-type repeat length. **G**. Southern blotting showing the absence of HRE in *C9orf72* in the edited lines. **H**. Immunostaining confirming expression of pluripotency markers in all cell lines used.

**Fig. S2. Quality control and exploratory analysis of RNA sequencing. A.** Panels showing expected (x axis) versus computed (y axis) expression for sequin spike-ins for both *C9orf72* HRE positive neurons (R^2=^0.74) and controls (R^2=^0.76). **B.** HOX gene expression in all samples, showing maximal expression for HOXA5, HOXB5 and HOXC4 indicating a dorsal hindbrain/cervical cord identity. **C.** Panel showing no expression of *PO5F1* (gene for Oct-3) and *NANOG*, as well as minimal expression of *KLF4*. **D.** Panel showing robust and similar expression of motor neuron markers Islet1, ChAT, NGFR and the limb transcription factor FOXP1. The MN precursor marker *OLIG2* and an early motor neuron marker *MNX1* (Hb9) are both lowly expressed.

**Fig. S3. Leading edge analysis reveals genes and pathways relevant to ALS.** Leading edge analysis enables an unbiased approach toward pathway discovery. All genes found in the leading edge of reactome pathways enriched in *C9orf72* HRE iPSMNs were selected (adjusted *p* < 0.05). Genes present in four or more pathways are plotted here. They include both AMPA and NMDA glutamate receptors, neurofilaments, endosomal pathway genes, synaptic genes and genes involved in calcium signalling. **B.** The KEGG ALS pathway is enriched in *C9orf72* HRE positive iPSMNs (adjusted *p* = 0.03), and genes driving this enrichment are presented here – there is a substantial overlap with the unbiased pathways presented above.

### Supplementary Tables and Sequences

**Supplementary Table 1.** Sources and dilutions of antibodies

| Antibody | Source | Secondary Antibody | Dilution | Application |
| --- | --- | --- | --- | --- |
| Islet-1 | DSHB | Alexaflour 568 goat Anti-Mouse | 1:200 | ICC |
| TUJ1 | Covance | Alexaflour 488 goat Anti-rabbit | 1:1000 | ICC |
| HB9 | DSHB | Alexaflour 568goat Anti-Mouse | 1:50 | ICC |
| Tuj-1 | Dako | HRP-conjugated anti-mouse | 1:1000 | WB |
| ChAT | Millipore | HRP-conjugated anti-mouse | 1:500 | WB |
| SSEA-4 | R&D | Alexaflour 633 Anti-Mouse | 1:50 | ICC |
| IgG3 (SSEA-4 isotype) | R&D | Alexaflour 633 Anti-Mouse | 1:50 | ICC |
| Oct4 | R&D | Alexaflour 488 Anti-Rat | 1:50 | ICC |
| IgG (Oct4 Isotype) | R&D | Alexaflour 488 Anti-Rat | 1:50 | ICC |
| Tra-1-60 | Biolegend | - | 1:100 | FC and ICC |
| IgM isotype | Biolegend | - | 1:100 | FC and ICC |
| Nanog | Cell Signaling | - | 1:150 | FC and ICC |
| IgG isotype | Cell Signaling | - | 1:150 | FC and ICC |

(ICC: Immunocytochemistry; WB: Western Blotting; FC: Flow Cytometry).

**Supplementary Table 2.**

Target sequences on *C9orf72* gene upstream of the repeat region, and oligonucleotides used to make the gRNA vectors.

| gRNA/Target site location | Target site sequence (5’>3’) PAM underlined | Oligo 1 (5’>3’) | Oligo 2 (5’>3’) |
| --- | --- | --- | --- |
| gRNA1 | AAGTAGTGGGGAGAGAGGGTGGG | CACCGAGTAGTGGGGAGAGAGGGT | AAACACCCTCTCTCCCCACTACTC |
| gRNA2 | GCTCTCACAGTACTCGCTGAGGG | CACCGCTCTCACAGTACTCGCTGA | AAACTCAGCGAGTACTGTGAGAGC |

**Supplementary Table 3.** Primer sequences.

| Primer name | Primer sequence (5’>3’) | Experiment |
| --- | --- | --- |
| SNP-F1  SNP-R1 | TATGGCCTGCCCAGAAGC  CGCGCGACTCCTGAGTTC | SNP detection, upstream of the repeat |
| SNP-F2  SNP-R2 | GTGGCGAGTGGGTGAGTG  CTTTCCACAACAGGAGCTGC | SNP detection, downstream of the repeat |
| HR-F  HR-R | TATGGCCTGCCCAGAAGC  CTTTCCACAACAGGAGCTGC | Donor template amplification |
| SDM-F  SDM-R | ATGCATGTGAACAAGAAAAGACCTGATAAAG  TCAGCGAGTACTGTGAGAGC | Site-directed mutagenesis |
| LoxP/Puro/tk-F  Puro-tk-r | CAACCGCAGCCTGTAGATAACTTCGTATAATGTATGCTATACGAAGTTATAGGAGTGGGAATTGGCTCC  CATAGAGCCCACCGCATCC | Gibson assembly experiment. |
| LoxP/Donor-F  Donor-R | TGCGGTGGGCTCTATGATAACTTCGTATAATGTATGCTATACGAAGTTATCAAGCTCTGGAACTCAGGAGTC  CTACAGGCTGCGGTTGTTTC | Gibson assembly experiment. |
| SEQ-F  SEQ-R | GTGGCGAGTGGGTGAGTG  CTTTCCACAACAGGAGCTGC | Direct sequencing. |
| Surv-F  Surv-R | ACTGCCTACCAAGCACAAACAA  GAAGGAGACAGCTCGGGTACT | Cleavage assays (Surveyor and T7E1) |
| RP1-F_ FAM  RP1-R (with G4C2 repeat)  RP1-Anchor | TGTAAAACGACGGCCAGTAGCTCTGGAACTCAGGAGTCGCG  CAGGAAACAGCTATGACCGGGCCCGCCCCGACCACGCCCCGGCCCCGGCCCCGG  CAGGAAACAGCTATGACC | Repeat-primed PCR (RP-PCR) |
| RP2-F (with G4C2 repeat)  RP2-Anchor  RP2-R_FAM | TACGCATCCCAGTTTGAGACGG4C2G4C2G4C2GGGG  TACGCATCCCAGTTTGAGACG  AGTCGCTAGAGGCGAAAGC | Repeat-primed PCR (RP-PCR) |
| Repeats-F  Repeats-R | AGCTCTGGAACTCAGGAGTCG  AGAAATGAGAGGGAAAGTAAAAATGC | Sequencing of repeat region |
| 5’-Out-F  EF1a-R | AGGTGAGTGGATTATGGGGTGG  ACTACCCCCGTCCGATTCTC | 5’ In-out |
| 5’SEQ-F  5’SEQ-R | CAAGTTCCGCCCACGTAAAA  TACAGGCTGCGGTTGTTTCC | Upstream Nsi1 |
| Puro-F  3’-Out-R | ATCCATGCCCACGCTACTG  CACCTTCTCCAACCTGGCTC | 3’In-out |
| 3’SEQ-R | CCCGGATGCAGGCAATT | Downstream Nsi1 site |
| Puro-R | ATAAACCCGCAGTAGCGTGGGCAT | Cassette excision |
| *Exc*-F  *Exc*-R | ACCGCAGCCTGTAGATAACTTC  AGTTCCAGAGCTTGATAACTTCGT | Cassette excision |
| *exon 2F*  *exon 3R* | CCCACTTCATAGAGTGTGTGTTG  TTCCATTCTCTCTGTGCCTTC | C9ORF72 ALL |
| *exon 4F*  *exon 5-UTR-R* | GAAATCACACAGTGTTCCTGAAGAA  ATCTGCTTCATCCAGCTTTTATGA | C9ORF72 V1 (SHORT) |
| *exon 8F*  *exon 9R* | CATGGCTCAGGATACGATCA  GGAAGGCTTTCACTAGAGTGTCTC | C9ORF72 V2+3 (LONG) |
| *exon 1AF*  *exon 2R* | GGGTCTAGCAAGAGCAGGTG  CGACATCACTGCATTCCAAC | C9ORF72 V3 (EX1A) |
| *Meth-F*  *Meth-R* | CAGTGTGAAAATCATGCTTGAGAGA  TTTGTGCTTGGTAGGCAGTG | Methylation |
| TOP1 | GTCCAAGCATAGCAACAGTGAAC  GCTCGAACCTTTTCCTCTTTTCG | qPCR validation |
| RPL13A | AGCTCATGAGGCTACGGAAAC  TTTATTGGGCTCAGACCAGGAG | qPCR validation |
| ATP5B | GCATTTGGGTGAGAGCACAG  TGGTGCACCAGAATCCAGTAC | qPCR validation |
| MAP13K13 | TGGCTATGAGAACCCCATGC  TTGGTCCTAACTGTGGCATCAG | qPCR validation |
| SYT11 | GACCTAGCTTTGATGTGTCACC  TGGTGGGTTCTTCTGCTTCTTC | qPCR validation |
| PCDHGB7 | TCATTGTCCAGCCACACAAG  CGGGGCTTGCTGTTTAAGAATC | qPCR validation |
| SV2A | CAACACGTTTTTCCGCAAC  TCAGACGGCTGTTCACAAAC | qPCR validation |
| ADAR | AGAAGGCAACTGCCCTACAG  CTTCTTTGCCTTTCCTGCAC | qPCR validation |
| TCEAL5 | AGAAGGCAACTGCCCTACAG  GTTCCCTTCTTTGCCTTTCC | qPCR validation |

**Supplementary Sequence 1.** Donor Template Sequence

pGEM-T Easy plasmid sequence

Right homology arm

Left homology arm

Exons 1a and 1b

Repeats region

gRNA regions

NsiI sites

LoxP sites

Ef1α Promoter

Puro/TK

BGH/P(A)

**Primer locations are underlined**

GGGCGAATTGGGCCCGACGTCGCATGCTCCCGGCCGCCATGGCGGCCGCGGGAATTCGATtatggcctgcccagaagctgatccagccatgcttcttgtacagcctgcagaactgtgagccattaaacttttctttataaattacccagtttcagttatttctttatagcagtgtaagaatggactaacacaattattaacgctagtcctcatgttgtacattaaatctctagatgtattagacgtaactgcaactttgtaccctaccctacaattttctttccccccaagccccccaaccaagggtctactctgtttctataaattcagttgttttttaattccacgtataagtgaagtacaactcagtgtagaaacttggtaaatgctagctacttgttataagctgtcagtcaaaataaaaatacagagatgaatctctaaattaagtgatttatttgggaagaaagaattgcaattagggcatacatgtagatcagatggtcttcggtatatccacacaacaaagaaaagggggaggttttgttaaaaaagagaaatgttacatagtgctctttgagaaaattcattggcactattaaggatctgaggagctggtgagtttcaactggtgagtgatggtggtagataaaattagagctgcagcaggtcattttagcaactattagataaaactggtctcaggtcacaacgggcagttgcagcagctggacttggagagaattacactgtgggagcagtgtcatttgtcctaagtgcttttctaccccctacccccactattttagttgggtataaaaagaatgacccaatttgtatgatcaactttcacaaagcatagaacagtaggaaaagggtctgtttctgcagaaggtgtagacgttgagagccattttgtgtatttattcctccctttcttcctcggtgaatgattaaaacgttctgtgtgatttttagtgatgaaaaagattaaatgctactcactgtagtaagtgccatctcacacttgcagatcaaaaggcacacagtttaaaaaacctttgtttttttacacatctgagtggtgtaaatgctactcatctgtagtaagtggaatctatacacctgcagaccaaaagacgcaaggtttcaaaaatctttgtgttttttacacatcaaacagaatggtacgtttttcaaaagttaaaaaaaaacaactcatccacatattgcaactagcaaatgacattccccagtgtgaaaatcatgcttgagagaattcttacatgtaaaggcaaaattgcgatgactttgcaggggaccgtgggattcccgcccgcagtgccggagctgtcccctaccagggtttgcagtggagttttgaatgcacttaacagtgtcttacggtaaaaacaaaatttcatccaccaattatgtgttgagcgcccactgcctaccaagcacaaacaaaaccattcaaaaccacgaaatcgtcttcactttctccagatccagcagcctcccctattaaggttcgcacacgctattgcgccaacgctcctccagagcgggtcttaagataaaagaacaggacaagttgccccgccccatttcgctagcctcgtgagaaaacgtcatcgcacatagaaaacagacagACGTAACCTACGGTGTCCCGCTAGGAAAGAGAGGTGCGTCAAACAGCGACAAGTTCCGCCCACGTAAAAGATGACGCTTGGTGTGTCAGCCGTCCCTGCTGCCCGGTTGCTTCTCTTTTGGGGGCGGGGTCTAGCAAGAGCAGGTGTGGGTTTAGGAGgtgtgtgtttttgtttttcccaccctctctccccactacttgctctcacagtactcgctgagATGCATGTGAACAAGAAAAGACCTGATAAAGATTAACCAGAAGAAAACAAGGAGGGAAACAACCGCAGCCTGTAGATAACTTCGTATAATGTATGCTATACGAAGTTATAGGAGTGGGAATTGGCTCCGGTGCCCGTCAGTGGGCAGAGCGCACATCGCCCACAGTCCCCGAGAAGTTGGGGGGAGGGGTCGGCAATTGAACCGGTGCCTAGAGAAGGTGGCGCGGGGTAAACTGGGAAAGTGATGTCGTGTACTGGCTCCGCCTTTTTCCCGAGGGTGGGGGAGAACCGTATATAAGTGCAGTAGTCGCCGTGAACGTTCTTTTTCGCAACGGGTTTGCCGCCAGAACACAGGTAAGTGCCGTGTGTGGTTCCCGCGGGCCTGGCCTCTTTACGGGTTATGGCCCTTGCGTGCCTTGAATTACTTCCACCTGGCTGCAGTACGTGATTCTTGATCCCGAGCTTCGGGTTGGAAGTGGGTGGGAGAGTTCGAGGCCTTGCGCTTAAGGAGCCCCTTCGCCTCGTGCTTGAGTTGAGGCCTGGCCTGGGCGCTGGGGCCGCCGCGTGCGAATCTGGTGGCACCTTCGCGCCTGTCTCGCTGCTTTCGATAAGTCTCTAGCCATTTAAAATTTTTGATGACCTGCTGCGACGCTTTTTTTCTGGCAAGATAGTCTTGTAAATGCGGGCCAAGATCTGCACACTGGTATTTCGGTTTTTGGGGCCGCGGGCGGCGACGGGGCCCGTGCGTCCCAGCGCACATGTTCGGCGAGGCGGGGCCTGCGAGCGCGGCCACCGAGAATCGGACGGGGGTAGTCTCAAGCTGGCCGGCCTGCTCTGGTGCCTGGCCTCGCGCCGCCGTGTATCGCCCCGCCCTGGGCGGCAAGGCTGGCCCGGTCGGCACCAGTTGCGTGAGCGGAAAGATGGCCGCTTCCCGGCCCTGCTGCAGGGAGCTCAAAATGGAGGACGCGGCGCTCGGGAGAGCGGGCGGGTGAGTCACCCACACAAAGGAAAAGGGCCTTTCCGTCCTCAGCCGTCGCTTCATGTGACTCCACGGAGTACCGGGCGCCGTCCAGGCACCTCGATTAGTTCTCGAGCTTTTGGAGTACGTCGTCTTTAGGTTGGGGGGAGGGGTTTTATGCGATGGAGTTTCCCCACACTGAGTGGGTGGAGACTGAAGTTAGGCCAGCTTGGCACTTGATGTAATTCTCCTTGGAATTTGCCCTTTTTGAGTTTGGATCTTGGTTCATTCTCAAGCCTCAGACAGTGGTTCAAAGTTTTTTTCTTCCATTTCAGGTGTCGTGAGGAATTTCGACTAGTGGCCACAACCATGGGGACCGAGTACAAGCCCACGGTGCGCCTCGCCACCCGCGACGACGTCCCCCGGGCCGTACGCACCCTCGCCGCCGCGTTCGCCGACTACCCCGCCACGCGCCACACCGTCGACCCGGACCGCCACATCGAGCGGGTCACCGAGCTGCAAGAACTCTTCCTCACGCGCGTCGGGCTCGACATCGGCAAGGTGTGGGTCGCGGACGACGGCGCCGCGGTGGCGGTCTGGACCACGCCGGAGAGCGTCGAAGCGGGGGCGGTGTTCGCCGAGATCGGCCCGCGCATGGCCGAGTTGAGCGGTTCCCGGCTGGCCGCGCAGCAACAGATGGAAGGCCTCCTGGCGCCGCACCGGCCCAAGGAGCCCGCGTGGTTCCTGGCCACCGTCGGCGTCTCGCCCGACCACCAGGGCAAGGGTCTGGGCAGCGCCGTCGTGCTCCCCGGAGTGGAGGCGGCCGAGCGCGCCGGGGTGCCCGCCTTCCTGGAGACCTCCGCGCCCCGCAACCTCCCCTTCTACGAGCGGCTCGGCTTCACCGTCACCGCCGACGTCGAGGTGCCCGAAGGACCGCGCACCTGGTGCATGACCCGCAAGCCCGGTGCCGGATCCATGCCCACGCTACTGCGGGTTTATATAGACGGTCCTCACGGGATGGGGAAAACCACCACCACGCAACTGCTGGTGGCCCTGGGTTCGCGCGACGATATCGTCTACGTACCCGAGCCGATGACTTACTGGCAGGTGCTGGGGGCTTCCGAGACAATCGCGAACATCTACACCACACAACACCGCCTCGACCAGGGTGAGATATCGGCCGGGGACGCGGCGGTGGTAATGACAAGCGCCCAGATAACAATGGGCATGCCTTATGCCGTGACCGACGCCGTTCTGGCTCCTCATATCGGGGGGGAGGCTGGGAGCTCACATGCCCCGCCCCCGGCCCTCACCCTCATCTTCGACCGCCATCCCATCGCCGCCCTCCTGTGCTACCCGGCCGCGCGATACCTTATGGGCAGCATGACCCCCCAGGCCGTGCTGGCGTTCGTGGCCCTCATCCCGCCGACCTTGCCCGGCACAAACATCGTGTTGGGGGCCCTTCCGGAGGACAGACACATCGACCGCCTGGCCAAACGCCAGCGCCCCGGCGAGCGGCTTGACCTGGCTATGCTGGCCGCGATTCGCCGCGTTTACGGGCTGCTTGCCAATACGGTGCGGTATCTGCAGGGCGGCGGGTCGTGGCGGGAGGATTGGGGACAGCTTTCGGGGACGGCCGTGCCGCCCCAGGGTGCCGAGCCCCAGAGCAACGCGGGCCCACGACCCCATATCGGGGACACGTTATTTACCCTGTTTCGGGCCCCCGAGTTGCTGGCCCCCAACGGCGACCTGTACAACGTGTTTGCCTGGGCCTTGGACGTCTTGGCCAAACGCCTCCGTCCCATGCACGTCTTTATCCTGGATTACGACCAATCGCCCGCCGGCTGCCGGGACGCCCTGCTGCAACTTACCTCCGGGATGGTCCAGACCCACGTCACCACCCCCGGCTCCATACCGACGATCTGCGACCTGGCGCGCACGTTTGCCCGGGAGATGGGGGAGGCTAACTGAGCTCTAGATGAAACGATATGGGCTGAATAACTAGAGCTCGCTGATCAGCCTCGACTGTGCCTTCTAGTTGCCAGCCATCTGTTGTTTGCCCCTCCCCCGTGCCTTCCTTGACCCTGGAAGGTGCCACTCCCACTGTCCTTTCCTAATAAAATGAGGAAATTGCATCGCATTGTCTGAGTAGGTGTCATTCTATTCTGGGGGGTGGGGTGGGGCAGGACAGCAAGGGGGAGGATTGGGAAGACAATAGCAGGCATGCTGGGGATGCGGTGGGCTCTATGATAACTTCGTATAATGTATGCTATACGAAGTTATcaagctctggaactcaggagtcgcgcgcta**ggggccggggccggggcc**ggggcgtggtcggggcgggcccgggggcgggcccggggcggggctgcggttgcggtgcctgcgcccgcggcggcggagGCGCAGGCGGTGGCGAGTGgtgagtgaATGCATaggcggcatcctggcgggtggctgtttggggttcggctgccgggarcgggtagaagcgggggctctcctcagagctcgacgcatttttactttccctctcatttctctgaccgaagctgggtgtcgggctttcgcctctagcgactggtggaattgcctgcatccgggccccgggcttcccggcggcggcggcggcggcggcggcgcagggacaagggatggggatctggcctcttccttgctttcccgccctcagtacccgagctgtctccttcccggggacccgctgggagcgctgccgctgcgggctcgagaaaagggagcctcgggtactgagaggcctcgcctgggggaaggccggagggtgggcggcgcgcggcttctgcggaccaagtcggggttcgctaggaacccgagacggtccctgccggcgaggagatcatgcgggatgagatgggggtgtggagacgcctgcacaatttcagcccaagcttctagagagtggtgatgacttgcatatgagggcagcaatgcaagtcggtgtgctccccattctgtgggacatgacctggttgcttcacagctccgagatgacacagacttgcttaaaggaagtgactattgtgacttgggcatcacttgactgatggtaatcagttgtctaaagaagtgcacagattacatgtccgtgtgctcattgggtctatctggccgcgttgaacaccaccaggctttgtattcagaaacaggagggaggtcctgcactttcccaggaggggtggccctttcagatgcaatcgagattgttaggctctgggagagtagttgcctggttgtggcagttggtaaatttctattcaaacagttgccatgcaccagttgttcacaacaagggtacgtaatctgtctggcattacttctacttttgtacaaaggatcaaaaaaaaaaaagatactgttaagatatgatttttctcagactttgggaaacttttaacataatctgtgaatatcacagaaacaagactatcatataggggatattaataacctggagtcagaatacttgaaatacggtgtcatttgacacgggcattgttgtcaccacctctgccaaggcctgccactttaggaaaaccctgaatcagttggaaactgctacatgctgatagtacatctgaaacaagaacgagagtaattaccacattccagattgttcactaagccagcatttacctgctccaggaaaaaattacaagcaccttatgaagttgataaaatattttgtttggctatgttggcactccacaatttgctttcagagaaacaaagtaaaccaaggaggacttctgtttttcaagtctgccctcgggttctattctacgttaattagatagttcccaggaggactaggttagcctacctattgtctgagaaacttggaactgtgagaaatggccagatagtgatatgaacttcaccttccagtcttccctgatgttgaagattgagaaagtgttgtgaactttctggtactgtaaacagttcactgtccttgaagtggtcctgggcagctcctgttgtggaaagATCACTAGTGAATTCGCGGCCGCCTGCAGGTCGACCATATGGGAGAGCTCCCAACGCGTTGGATGCATAGCTTGAGTATTCTATAGTGTCACCTAAATAGCTTGGCGTAATCATGGTCATAGCTGTTTCCTGTGTGAAATTGTTATCCGCTCACAATTCCACACAACATACGAGCCGGAAGCATAAAGTGTAAAGCCTGGGGTGCCTAATGAGTGAGCTAACTCACATTAATTGCGTTGCGCTCACTGCCCGCTTTCCAGTCGGGAAACCTGTCGTGCCAGCTGCATTAATGAATCGGCCAACGCGCGGGGAGAGGCGGTTTGCGTATTGGGCGCTCTTCCGCTTCCTCGCTCACTGACTCGCTGCGCTCGGTCGTTCGGCTGCGGCGAGCGGTATCAGCTCACTCAAAGGCGGTAATACGGTTATCCACAGAATCAGGGGATAACGCAGGAAAGAACATGTGAGCAAAAGGCCAGCAAAAGGCCAGGAACCGTAAAAAGGCCGCGTTGCTGGCGTTTTTCCATAGGCTCCGCCCCCCTGACGAGCATCACAAAAATCGACGCTCAAGTCAGAGGTGGCGAAACCCGACAGGACTATAAAGATACCAGGCGTTTCCCCCTGGAAGCTCCCTCGTGCGCTCTCCTGTTCCGACCCTGCCGCTTACCGGATACCTGTCCGCCTTTCTCCCTTCGGGAAGCGTGGCGCTTTCTCATAGCTCACGCTGTAGGTATCTCAGTTCGGTGTAGGTCGTTCGCTCCAAGCTGGGCTGTGTGCACGAACCCCCCGTTCAGCCCGACCGCTGCGCCTTATCCGGTAACTATCGTCTTGAGTCCAACCCGGTAAGACACGACTTATCGCCACTGGCAGCAGCCACTGGTAACAGGATTAGCAGAGCGAGGTATGTAGGCGGTGCTACAGAGTTCTTGAAGTGGTGGCCTAACTACGGCTACACTAGAAGAACAGTATTTGGTATCTGCGCTCTGCTGAAGCCAGTTACCTTCGGAAAAAGAGTTGGTAGCTCTTGATCCGGCAAACAAACCACCGCTGGTAGCGGTGGTTTTTTTGTTTGCAAGCAGCAGATTACGCGCAGAAAAAAAGGATCTCAAGAAGATCCTTTGATCTTTTCTACGGGGTCTGACGCTCAGTGGAACGAAAACTCACGTTAAGGGATTTTGGTCATGAGATTATCAAAAAGGATCTTCACCTAGATCCTTTTAAATTAAAAATGAAGTTTTAAATCAATCTAAAGTATATATGAGTAAACTTGGTCTGACAGTTACCAATGCTTAATCAGTGAGGCACCTATCTCAGCGATCTGTCTATTTCGTTCATCCATAGTTGCCTGACTCCCCGTCGTGTAGATAACTACGATACGGGAGGGCTTACCATCTGGCCCCAGTGCTGCAATGATACCGCGAGACCCACGCTCACCGGCTCCAGATTTATCAGCAATAAACCAGCCAGCCGGAAGGGCCGAGCGCAGAAGTGGTCCTGCAACTTTATCCGCCTCCATCCAGTCTATTAATTGTTGCCGGGAAGCTAGAGTAAGTAGTTCGCCAGTTAATAGTTTGCGCAACGTTGTTGCCATTGCTACAGGCATCGTGGTGTCACGCTCGTCGTTTGGTATGGCTTCATTCAGCTCCGGTTCCCAACGATCAAGGCGAGTTACATGATCCCCCATGTTGTGCAAAAAAGCGGTTAGCTCCTTCGGTCCTCCGATCGTTGTCAGAAGTAAGTTGGCCGCAGTGTTATCACTCATGGTTATGGCAGCACTGCATAATTCTCTTACTGTCATGCCATCCGTAAGATGCTTTTCTGTGACTGGTGAGTACTCAACCAAGTCATTCTGAGAATAGTGTATGCGGCGACCGAGTTGCTCTTGCCCGGCGTCAATACGGGATAATACCGCGCCACATAGCAGAACTTTAAAAGTGCTCATCATTGGAAAACGTTCTTCGGGGCGAAAACTCTCAAGGATCTTACCGCTGTTGAGATCCAGTTCGATGTAACCCACTCGTGCACCCAACTGATCTTCAGCATCTTTTACTTTCACCAGCGTTTCTGGGTGAGCAAAAACAGGAAGGCAAAATGCCGCAAAAAAGGGAATAAGGGCGACACGGAAATGTTGAATACTCATACTCTTCCTTTTTCAATATTATTGAAGCATTTATCAGGGTTATTGTCTCATGAGCGGATACATATTTGAATGTATTTAGAAAAATAAACAAATAGGGGTTCCGCGCACATTTCCCCGAAAAGTGCCACCTGATGCGGTGTGAAATACCGCACAGATGCGTAAGGAGAAAATACCGCATCAGGAAATTGTAAGCGTTAATATTTTGTTAAAATTCGCGTTAAATTTTTGTTAAATCAGCTCATTTTTTAACCAATAGGCCGAAATCGGCAAAATCCCTTATAAATCAAAAGAATAGACCGAGATAGGGTTGAGTGTTGTTCCAGTTTGGAACAAGAGTCCACTATTAAAGAACGTGGACTCCAACGTCAAAGGGCGAAAAACCGTCTATCAGGGCGATGGCCCACTACGTGAACCATCACCCTAATCAAGTTTTTTGGGGTCGAGGTGCCGTAAAGCACTAAATCGGAACCCTAAAGGGAGCCCCCGATTTAGAGCTTGACGGGGAAAGCCGGCGAACGTGGCGAGAAAGGAAGGGAAGAAAGCGAAAGGAGCGGGCGCTAGGGCGCTGGCAAGTGTAGCGGTCACGCTGCGCGTAACCACCACACCCGCCGCGCTTAATGCGCCGCTACAGGGCGCGTCCATTCGCCATTCAGGCTGCGCAACTGTTGGGAAGGGCGATCGGTGCGGGCCTCTTCGCTATTACGCCAGCTGGCGAAAGGGGGATGTGCTGCAAGGCGATTAAGTTGGGTAACGCCAGGGTTTTCCCAGTCACGACGTTGTAAAACGACGGCCAGTGAATTGTAATACGACTCACTATA
